# Cell-to-cell variability in JAK2/STAT5 pathway components and cytoplasmic volumes define survival threshold in erythroid progenitor cells

**DOI:** 10.1101/866871

**Authors:** Lorenz Adlung, Paul Stapor, Christian Tönsing, Leonard Schmiester, Luisa E. Schwarzmüller, Dantong Wang, Jens Timmer, Ursula Klingmüller, Jan Hasenauer, Marcel Schilling

**Affiliations:** Division Systems Biology of Signal Transduction, German Cancer Research Center (DKFZ), 69120 Heidelberg, Germany; Helmholtz Zentrum München - German Research Center for Environmental Health, Institute of Computational Biology, 85764 Neuherberg, Germany; Technische Universität München, Center for Mathematics, Chair of Mathematical Modeling of Biological Systems, 85748 Garching, Germany; Institute of Physics, University of Freiburg, 79104 Freiburg, Germany; CIBSS - Centre for Integrative Biological Signalling Studies, University of Freiburg, 79104 Freiburg, Germany; Translational Lung Research Center (TLRC), Member of the German Center for Lung Research (DZL), 69120 Heidelberg, Germany; Faculty of Mathematics and Natural Sciences, University of Bonn, 53113 Bonn, Germany

**Keywords:** single-cell modeling, JAK/STAT, signal transduction, Epo

## Abstract

Survival or apoptosis is a binary decision in individual cells. Yet, at the cell population level, a graded increase in survival of CFU-E cells is observed upon stimulation with Erythropoietin (Epo). To identify components of JAK2/STAT5 signal transduction that contribute to the graded population response, a cell population-level model calibrated with experimental data was extended to study the behavior in single cells. The single-cell model showed that the high cell-to-cell variability in nuclear phosphorylated STAT5 is caused by variability in the amount of EpoR:JAK2 complexes and of SHP1 as well as the extent of nuclear import due to the large variance in the cytoplasmic volume of CFU-E cells. 24 to 118 pSTAT5 molecules in the nucleus for 120 min are sufficient to ensure cell survival. Thus, variability in membrane-associated processes are responsible to convert a switch-like behavior at the single-cell level to a graded population level response.

**Highlights:**

- Mathematical modeling enables integration of heterogeneous data
- Single-cell modeling captures binary decision process
- Multiple sources of cell-to-cell variability in erythroid progenitor cells
- Minimal amount of active STAT5 sufficient for survival of erythroid progenitor cells

## Introduction

Signal transduction has been intensively studied in the last decades at the cell population level utilizing immunoblotting and bulk gene-expression analyses. However, information processing is highly dynamic and occurs at the single-cell level with considerable cell-to-cell variability (Taniguchi et al, 2010). This variability can be beneficial in some contexts and harmful in others (Raj & van Oudenaarden, 2008). For example, cell-to-cell variability can improve robustness of signal transduction responses (Paszek et al, 2010) but also lead to incomplete growth inhibition of tumor cells (Niepel et al, 2009). It has been reported that while cell-to-cell variability is fundamental to most molecular processes in cells, quantitative assessment of how this influences information processing through intracellular signal transduction networks is almost completely lacking (Pelkmans, 2012). Furthermore, several cellular responses such as apoptosis, cell cycle entry or epithelial to mesenchymal transition are binary cell fate decisions in single cells, but typically appear to occur as a graded response at the population level due to cell-to-cell variability.

For the adequate production of mature erythrocytes, which is essential to maintaining constant plasma concentrations of hemoglobin, the survival of erythroid progenitor cells at the colony forming erythroid stage (CFU-E) has to be tightly controlled (Nijhof & Wierenga, 1983). A homogenous switch-like induction of survival in all CFU-E cells by the hormone erythropoietin (Epo), a key regulator of erythropoiesis (Richmond et al, 2005), would lead to detrimental fluctuations in hemoglobin levels. Therefore, a graded input-output relationship of survival of CFU-E cells as a function of Epo concentrations is beneficial (Koulnis et al, 2014). Yet, since in individual CFU-E cells Epo-induced signal transduction that promotes cell survival triggers a binary decision, to achieve a graded output at the population level, heterogeneity in the sensitivity of CFU-E cells to Epo is required.

In CFU-E cells, the JAK2/STAT5 signal transduction pathway is a key mediator of cell survival (Socolovsky et al, 2001). It facilitates rapid signal transduction and connects binding of Epo to its cell surface receptor, the EpoR, with gene expression in the nucleus (Swameye et al, 2003) and cell survival (Bachmann et al, 2011). The Epo-EpoR complex activates the receptor-associated Janus kinase JAK2, which phosphorylates the cytoplasmic tail of the EpoR on multiple tyrosine residues (Klingmüller et al, 1996). STAT5 is recruited to the phosphorylated receptor complex and is in turn phosphorylated by JAK2 (Gouilleux et al, 1995). Phosphorylated STAT5 (pSTAT5) molecules form dimers (Boehm et al, 2014), translocate to the nucleus and induce the expression of anti-apoptotic target genes, e.g. *Bcl2l1* for immediate control of cell survival (Socolovsky et al, 1999). Other STAT5 target genes include *Cish* (Yoshimura et al, 1995) and *Socs3* (Sasaki et al, 2000) that encode the negative feedback regulators cytokine-inducible SH2 domain containing protein (CIS) and suppressor of cytokine signaling 3 (SOCS3), which attenuate signal transduction. The SH2 domain-containing protein tyrosine phosphatase 1 (SHP1) is a cytosolic protein that is recruited to the activated receptor via its SH2 domain (Neel et al, 2003). Receptor recruitment activates the phosphatase activity of SHP1 and thereby causes dephosphorylation of JAK2 leading to termination of signal transduction (Klingmüller et al, 1996). Since the activation of the core JAK2/STAT5 signal transduction cascade is controlled by multiple transcription-induced feedbacks and an induced activity feedback, the reaction network coupling Epo to survival of CFU-E cells are non-linear. This renders the outcome difficult to predict.

Dynamic pathway models provide suitable tools to identify systems properties in such non-linear reaction networks. Previously, we established a dynamic pathway model of the Epo-induced JAK2/STAT5 signal transduction pathway in CFU-E cells and calibrated the model parameters using population-average data (Bachmann et al, 2011). This population average model of the JAK2/STAT5 pathway enabled us to discern that the activation of the pathway is terminated by two dose-dependent transcriptional feedback regulators, SOCS3 and CIS, which show maximal expression after approximately 60 min of Epo stimulation and operate as noise filters in the pathway. Further, the model showed that the integral of pSTAT5 in the nucleus observed within 60 min of Epo stimulation correlates with the extent of survival of CFU-E cells (Bachmann et al, 2011). However, this mathematical model only described the average activation of the pathway in a population of CFU-E cells without taking cell-to-cell variability into account. Therefore, it remained elusive how the gradual Epo-induced increase in STAT5 phosphorylation and survival of CFU-E cells at the cell population level relates to the switch-like all-or-none survival decision that occurred in an individual cell.

The elucidation of mechanisms that lead to cell-to-cell variability in responses requires mechanistic modeling approaches capturing properties in individual cells. While computational frameworks for the quantitative dynamical modeling of population-level data are well established (Balsa-Canto & Banga, 2011; Hoops et al, 2006; Raue et al, 2015), mechanistic modeling of single-cell data remains challenging. Available approaches for the mechanistic description of cell-to-cell variability include stochastic modeling for gene expression (Neuert et al, 2013), mixed-effect modeling for signal transduction (Karlsson et al, 2015) and a variety of hybrid approaches (Fröhlich et al, 2018; Loos et al, 2018; Toni & Tidor, 2013; Zechner et al, 2012). However, these mathematical modeling techniques are computationally very demanding, which renders model establishment and comparison challenging. Therefore, it was of importance to establish a method that enables efficient and yet accurate parameter estimation in single-cell models. For the calibration of such mathematical models, appropriate quantitative experimental data is required. Recently, single-cell RNA sequencing approaches have been utilized (Buettner et al, 2015; Dalerba et al, 2011) to study gene expression in single cells. Clustering algorithms facilitated the identification of distinct cell populations based on these data sets. While these data could also be employed to analyze cell-to-cell variability in a specific cell population, usually only a few hundred unique transcripts are sequenced per cell, limiting the resolution of these methods. More importantly, since it has been shown that the correlation between mRNA expression levels and protein abundance is low (Schwanhausser et al, 2011) and intracellular information processing, which links e.g. changes in ligand concentration to cell fate decisions, is executed by complex non-linear reaction networks, it is essential to assess alterations of proteins at the single-cell level. While single-cell time-lapse microscopy allows to follow changes in proteins in individual cells over time, options for monitoring signal transduction components and multiplexing are limited in this technique because only few fluorophores for simultaneous quantification are available and the required tagging with fluorophores might change the properties of proteins (Weill et al, 2019). In contrast, flow cytometry and mass cytometry enable snapshot measurements of dozens of markers in individual cells (Bodenmiller et al, 2012; Perfetto et al, 2004).

Studies combining experimental data and mathematical modeling previously investigated molecular mechanisms for cell fate decisions. Several studies suggested that if the abundance of certain components exceeds a limit, a cellular decision is taken suggesting the existence of an absolute threshold. For example, the absolute abundance of the spindle assembly checkpoint proteins were reported to be crucial for mitosis in fission yeast. Because even small variations in checkpoint protein abundance can strongly impact spindle assembly checkpoint signaling, the level of the relevant checkpoint proteins was shown to be kept within a narrow window by stoichiometric inhibition (Heinrich et al, 2013). Likewise, for apoptosis mediated by TRAIL in human cell lines, it was demonstrated that naturally occurring differences in protein abundances are the primary causes of cell-to-cell variability in the timing and probability of death. If the cellular abundance of tBid reaches a certain value, an irreversible process mediated by positive feedback loops is triggered leading to apoptosis (Spencer et al, 2009). However, other studies showed that for some cellular decisions, a fold-change can be decisive suggesting a relative threshold. For example, it was shown in the Wnt signal transduction pathway that gene expression and the embryonic phenotype correlated with the fold-change in β-catenin abundance (post stimulation/pre-stimulation), rather than the absolute abundance (Goentoro & Kirschner, 2009). Similarly, in the EGF signal transduction pathway, H1299 human non-small cell lung cancer cells showed a high cell-to-cell variability in ERK2 nuclear abundance. Evidence was provided that the amount of ERK2 entering the nucleus upon EGF stimulation is proportional to the basal level of nuclear ERK2 in each cell, suggesting a fold-change response mechanism (Cohen-Saidon et al, 2009). Our previous studies on Epo-induced survival of CFU-E cells showed that at the cell population level the integration of the amount of nuclear pSTAT5 for 60 min after Epo-stimulation correlated with survival of CFU-E cells suggesting that cell survival is an early decision (Bachmann et al, 2011). However, it remained unresolved whether in an individual CFU-E cell an absolute or relative threshold of pSTAT5 in the nucleus is decisive for Epo-induced cell survival.

Here we report the development of a population average model of the Epo-induced JAK2/STAT5 signal transduction pathway in primary murine erythroid progenitor cells that captures cellular population-average data. This model in combination with flow cytometric analyses of STAT5 and pSTAT5 is utilized to create a mixed-effect model of pathway activation at the single-cell level. Analysis of the single-cell model suggests that a high variability in the amounts of the EpoR:JAK2 complex and the phosphatase SHP1 and in the nuclear translocation rates of STAT5 contribute to the detected high variability in nuclear pSTAT5. With this approach we identify a relative threshold of nuclear pSTAT5 deciding survival in CFU-E cells and elucidate the mechanisms converting the switch-like survival decision in individual CFU-E cells to a graded response at the population level.

## Results

### Cell-to-cell variability of phosphorylated STAT5 in primary erythroid progenitor cells

As previously reported, the application of an increasing dose of erythropoietin (Epo) results in a graded increase of surviving CFU-E erythroid progenitor cells (Bachmann et al, 2011) (Figure S1). The key intracellular integrator of Epo-induced survival signal transduction is the latent transcription factor STAT5 that is activated by tyrosine phosphorylation (pSTAT5). We hypothesized that variability in the expression level of total STAT5 in individual cells causes variability in pSTAT5 and explains that a binary decision in individual cells is converted to a graded response at the cell population level.

To experimentally evaluate the expression of STAT5 in CFU-E cells, we stimulated the cells for 15 min with a broad range of Epo doses that support cell survival (Figure S1) and monitored the expression of total STAT5 and of Epo-induced phosphorylation of STAT5 by flow cytometry (Figure 1A-E). In this study we used -similar to most studies in the field (Herzenberg et al, 2006; Moore & Parks, 2012; Parks et al, 2006) -the logicle scale for the visualization and analysis of the flow cytometry data. The logicle scale allows to deal with the negative values resulting from signal compensation.

**Figure 1:**
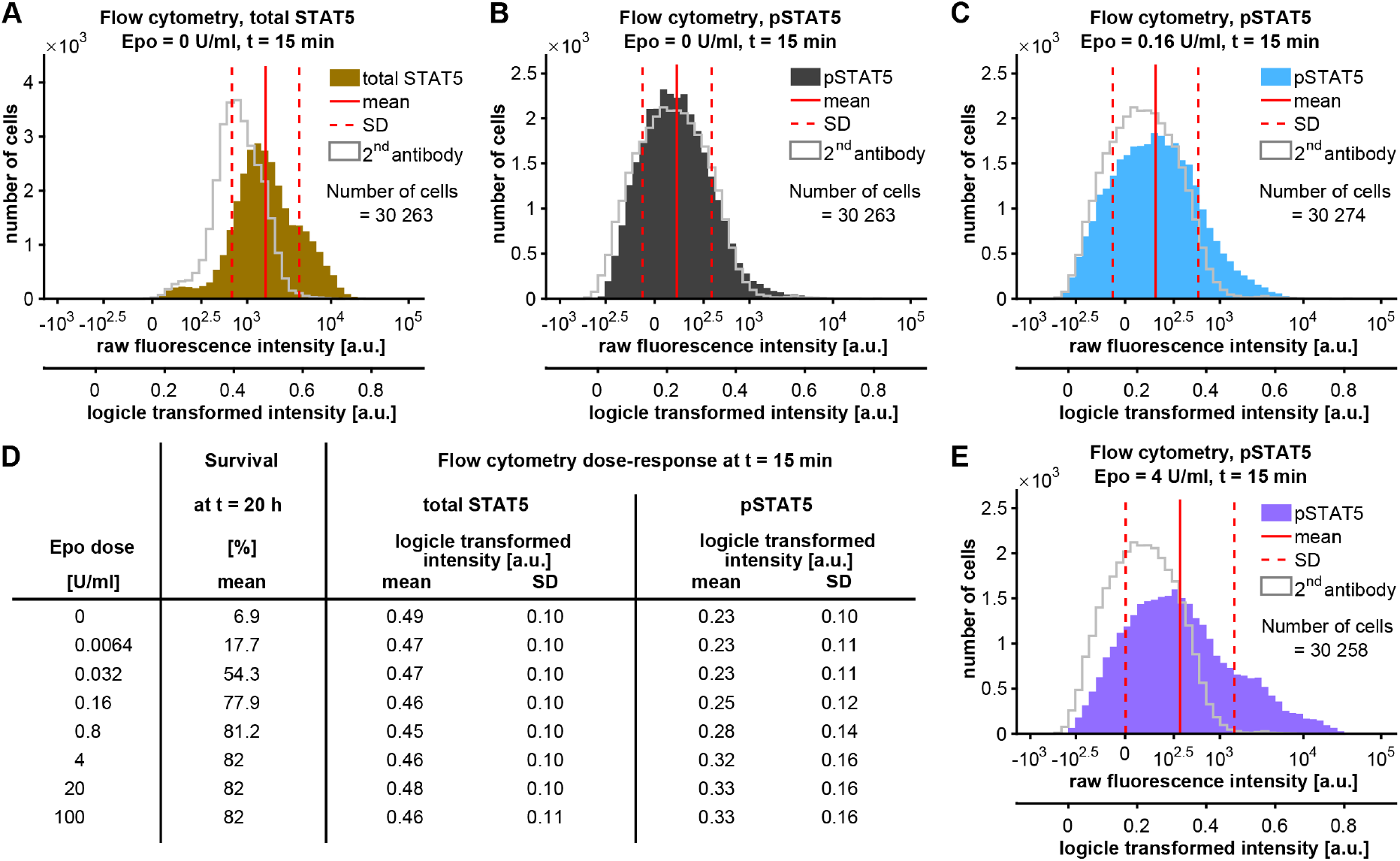
Cell-to-cell variability in total and phosphorylated STAT5. (A) Unstimulated growth factor-depleted CFU-E cells were fixed, permeabilized and intracellularly stained with goat anti-STAT5 primary antibody and anti-goat FITC secondary antibody (brown) or only the secondary antibody (grey line). Growth factor-depleted CFU-E cells were (B) unstimulated or were stimulated with (C) 0.16 U/ml Epo or (E) 4 U/ml Epo for 15 min and were subjected to intracellular staining with rabbit anti-pSTAT5 antibody and with anti-rabbit APC secondary antibody – dark grey (unstimulated), blue (0.16 U/ml Epo), purple (4 U/ml Epo) – or secondary antibody only (grey line). Raw fluorescence intensities are plotted in a bi-exponential manner and in each panel the corresponding logicle scale is indicated. (D) Growth factor-depleted CFU-E cells were stimulated for 20 h with the indicated Epo concentrations and the corresponding fraction of surviving CFU-E cells was extrapolated (Figure S1) based on previously published data (Bachmann et al, 2011). For each of these Epo concentrations the extracted mean and standard deviation (SD) of flow cytometric measurements of total STAT5 and pSTAT5 transformed to the logicle scale are displayed.

As shown in Figure 1A, in unstimulated CFU-E cells, total STAT5 was detected with a mean fluorescence intensity of 0.49 on the logicle scale and a standard deviation (SD) of 0.10. Intracellular staining with the secondary antibody alone revealed that a substantial fraction of the detected signal was due to non-specific binding of the secondary antibody. Upon stimulation with increasing doses of Epo the mean fluorescent intensity of total STAT5 varied between 0.45 and 0.48 and showed an SD of 0.10 to 0.11 and was thus rather unaffected by Epo. The flow cytometric detection of pSTAT5 yielded in the unstimulated situation a mean fluorescence intensity of 0.23 and a SD of 0.10. The detected signal was to a large extent overlapping with the signal generated by an incubation with the secondary antibody alone (Figure 1B). However, upon stimulation with 0.16 U/ml Epo, the mean fluorescence intensity measured for pSTAT5 shifted to a higher mean value (0.25) and was distributed more broadly (SD of 0.12) (Figure 1C). This effect was even more pronounced upon stimulation with 4 U/ml Epo, with a mean fluorescence intensity of 0.32 and a SD of 0.16 (Figure 1E), which remained the same upon adding even higher Epo doses (Figure 1D). While the fraction of surviving cells for the respective Epo concentration as interpolated from our previously reported data (Bachmann et al, 2011) increased in a graded fashion (Figure S1), the mean fluorescence intensities and SD of total STAT5 did not change with increasing Epo doses. On the contrary, for pSTAT5 we observed a gradual increase in the mean fluorescence intensity that saturated similar to the Epo-induced cell survival at a concentration of 4 U/ml Epo. Interestingly, the SD observed for pSTAT5 was considerably larger than for total STAT5 and showed an increase in response to rising Epo doses that correlated with the survival responses (Figure 1D).

Thus, we concluded that cell-to-cell variability in the expression of total STAT5 alone was not sufficient to explain the Epo dose-dependent increase in the variance of the key integrator of survival signaling pSTAT5. Rather, variability in components and non-linear reactions controlling the formation of pSTAT5 might have a major impact on survival decisions in individual cells.

### Dynamical modeling of Epo-induced JAK2/STAT5 signal transduction at the population level

To identify pathway components and reaction rates in the JAK2/STAT5 signaling pathway that vary from cell to cell and could cause the observed variability in STAT5 phosphorylation, a strategy had to be developed that permits an in-depth analysis of cell-to-cell variability by mechanistic modeling.

First, we established a mechanistic mathematical model which describes the system on a population-average level utilizing our previously published mathematical model of the JAK2/STAT5 signaling pathway. This model consists of coupled ordinary differential equations (ODE) and describes Epo-induced JAK2/STAT5 signaling and cell survival at the cell population level (Bachmann et al, 2011) (Figure 2A). So far, calibration of model parameters in dynamic pathway models was performed using quantitative immunoblotting data, quantitative mass spectrometry and qRT-PCR data that assess the average dynamic behavior of the cell population. However, flow cytometry offers the advantage to determine the average behavior of the cell population by calculating the mean fluorescence intensities, but also provides information on the distribution of the fluorescence intensity across the cell population and thus enables a link to the single-cell level. Therefore, to calibrate the parameters of our population average model, we collected quantitative data acquired in CFU-E cells with a wide variety of experimental approaches and conditions: (i) Quantitative immunoblotting analysis of total cell lysates for pSTAT5 and tSTAT5 and of cytoplasmic lysates for pEpoR, pJAK2, CIS, SOCS3 and SHP1 served as a readout to determine the average Epo-induced activation of the JAK2/STAT5 pathway (Figure 2B and S2 B5, B7, C5-C6, D1-D4, D7-E4, E6-F2, F5-G4, G6, H7, K1-K4, K6-K7). (ii) Targeted mass spectrometry using one-source peptide/phosphopeptide ratio standards (Boehm et al, 2014) allowed to determine relative average amounts of pSTAT5 in response to Epo stimulation (Figure 2C and S2 L1, L2). (iii) Cell fractionation experiments, separating the cytoplasmic and the nuclear compartment, provided access to the average dynamics of pSTAT5 localization (Figure 2D-E and S2 B1-B4, B6, C1-C4, C7, D5, D6, E5, F3, F4, G5, G7, H6, K5). (iv) qRT-PCR experiments (Figure S2 I5-I7, J1-J7) revealed the induction dynamics of the transcriptional feedbacks. (v) The mean fluorescence intensities of the flow cytometry data on pSTAT5 provided time and Epo-dose resolved information on the average pSTAT5 concentration in the cell population (Figure 2F and S2 A1-A3, B7). Altogether a curated data set of 516 published data points (Bachmann et al, 2011) and 469 newly acquired data points providing information on the average dynamic behavior of the JAK2/STAT5 pathway in the cell population was assembled (Figure 2G). A major challenge for the calibration of the population-average model was the use of data with such different scaling and error structures, e.g. immunoblotting data and flow cytometry data. It turned out that it is crucial to simultaneously estimate model parameters and the measurement noise (Raue et al, 2013) on the scale given by the measurement technique: The measured values are given for immunoblotting and qRT-PCR measurements on a log_10_ scale (Kreutz et al, 2007), for mass spectrometry as a percentage of pSTAT5 relative to total STAT5 (Boehm et al, 2014) and for flow cytometry data on a logicle scale (Moore & Parks, 2012). For the estimation of the model parameters based on these data sets we used a multi-start local optimization. Additionally, we reduced the complexity of the model without substantially impairing the model agreement to the comprehensive data set. We applied an iterative method, which relies on profile likelihood calculation (Maiwald et al, 2016). The resulting population-average model (Figure 2A) differed from the previously published model in three aspects: (i) SHP1 activation was simplified by considering the total amount of SHP1 as not-limiting, (ii) SOCS3 transcription and translation process were summarized in a single reaction and (iii) the CIS transcriptional delay was shortened, which was modeled by the linear chain trick (MacDonald, 1976) (for details and biological interpretation, see STAR Methods section “Model reduction”). Calibrating the reduced model to the experimental data showed a favorable convergence (Raue et al, 2013) during parameter optimization (Figure 2H), a good agreement (see Figure S2) and a high correlation (*ρ*=0.978, see Figure S4A, B) of experimental data and model output for the best fit. The profile likelihood calculation revealed that 19 out of 21 of the estimated dynamic parameters of the population-average model are practically identifiable at a confidence level of 95%, and all 21 parameters are practically identifiable at a confidence level of 68% (Figure S3). Accordingly, the calibrated population-average model accurately described the experimental data (Figure S2). For example, the mathematical model was capable to capture the dynamics of pSTAT5 upon stimulation of CFU-E cells with 5 U/ml Epo such as the rapid activation dynamics with a peak of absolute pSTAT5 at 33 min measured by quantitative immunoblotting in the whole cell lysates (Figure 2B) and the sharp peak of 67% phosphorylation of cytoplasmic STAT5 at 23 min returning to a steady state level of 24% after 120 min as determined by quantitative mass spectrometry (Figure 2C). Furthermore, by cell fractionation followed by immunoblotting analysis, a peak of absolute pSTAT5 was detected at 13 min after stimulation with 5 U/ml Epo in the cytoplasmic lysates of CFU-E cells (Figure 2D), while nuclear pSTAT5 was rather sustained (Figure 2E) and both features were faithfully represented by the mathematical model. Likewise, the mathematical model was able to describe the Epo dose-dependent shift of the peak of STAT5 phosphorylation in CFU-E cells to earlier time points with a peak at 82 min for 0.032 U/ml Epo, a peak at 46 min for 0.8 U/ml Epo and a peak at 28 min for 20 U/ml Epo, as measured by flow cytometry (Figure 2F). Finally, the mathematical model was also able to describe dynamics of STAT5 phosphorylation detected for the stimulation with additional Epo doses (Figure S2).

**Figure 2:**
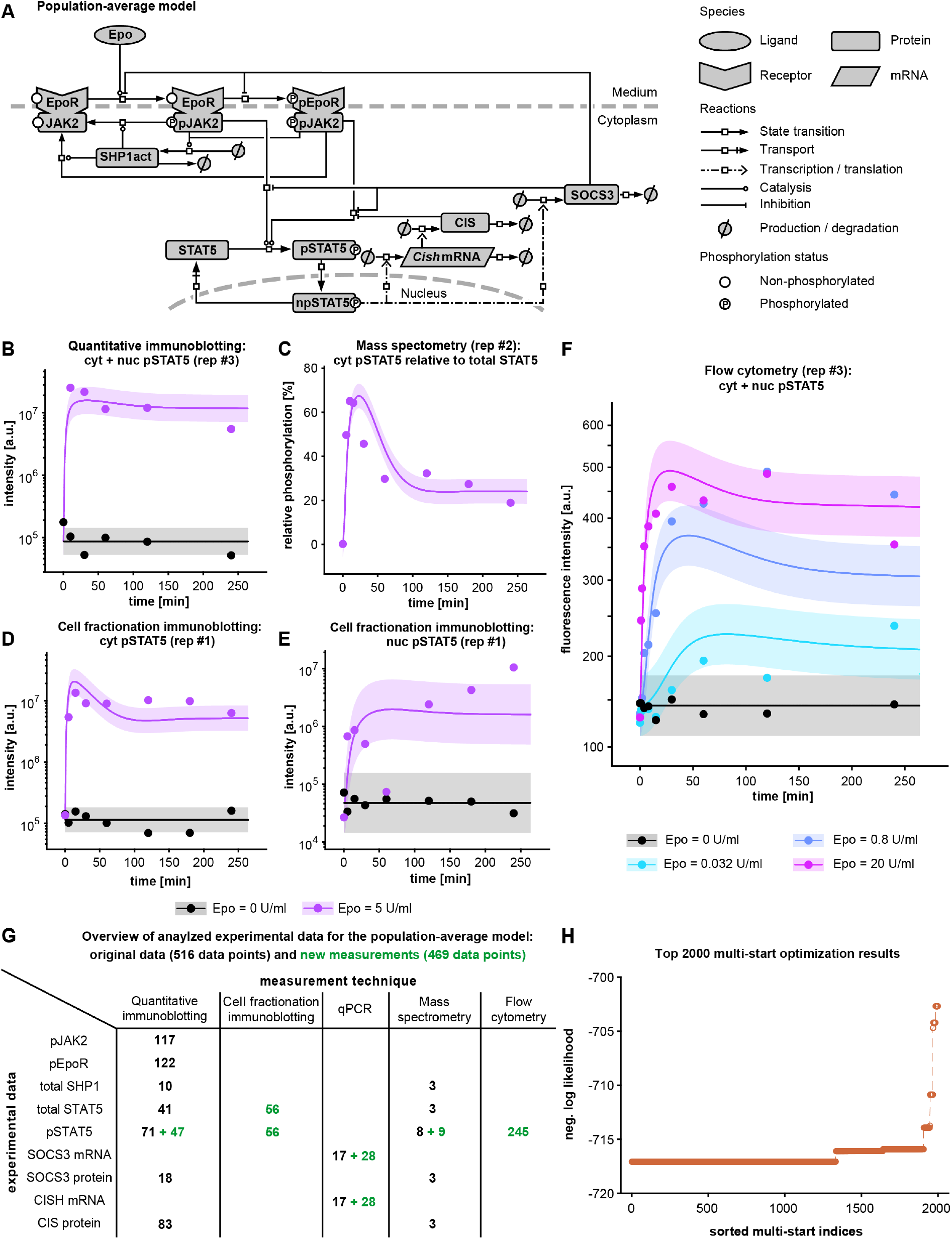
Modeling of population-average dynamics. (A) The structure of the population-average JAK2/STAT5 model is represented by a process diagram displayed according to Systems Biology Graphical Notation (Le Novere et al, 2009). (B-F) Illustrative experimental data (filled circles) and model simulations (solid lines) for pSTAT5 (see Figure S2 for complete data set with full range of Epo doses, with all replicates and observed model outputs). Data shown were generated by (B) quantitative immunoblotting from whole-cell lysates, (C) quantitative mass spectrometry from cytoplasmic lysates using one-source peptide/phosphopeptide ratio standards, (D,E) quantitative immunoblotting from cytoplasmic and nuclear lysates upon cell fractionation experiments and (F) population-average fluorescence intensities of pSTAT5 measured by flow cytometry (see Figure S2, A1-3 for all Epo doses). (G) Summary of experimental data used for calibration of the population-based mathematical model. Number of data measurements in published dataset (Bachmann et al, 2011) are given in black, new measurements of this study are given in green. (H) Multi-start optimization results, shown as likelihood waterfall plot (Raue et al, 2013). The best 2000 of 5000 local optimization runs are shown, plateaus indicate convergence to local or global optima. See Figure S3 for profile likelihood of estimated model parameters and Figure S4A, B for correlation of data and model output.

To conclude, we established a calibrated population-average model for the JAK2/STAT5 signal transduction pathway in CFU-E cells that is capable to accurately describe the dynamics of pathway activation in these cells including the average level of pSTAT5 measured by flow cytometry, which thereby forms the basis for the analysis of the pathway at the single-cell level.

### Establishment of parameter estimation methods to model Epo-induced JAK2/STAT5 signal transduction in single cells

To describe Epo-induced JAK2/STAT5 signal transduction at the single-cell level, we compared different approximation methods to enable parameter estimation in single-cell models. A faithful representation of the cell-to-cell variability at a given time point in a cell population had to be balanced with computational costs of the utilized multi-start parameter estimation. Because the number of molecules ranged from around 1000 molecules for the EpoR to around 20 000 molecules per cell for STAT5 (Bachmann et al, 2011) and was therefore sufficiently large, we employed a deterministic mixed-effect modeling approach (Fröhlich et al, 2017). Mixed-effect modeling allowed each parameter to be either fixed or variable across cells. Parameters, which are assumed to be variable across cells, were referred to as “random effects”, otherwise they were called “fixed effects”. Random effects should follow a multivariate normal distribution, which was parametrized by the population mean and a covariance matrix. Hence, compared to the population-average JAK2/STAT5 model, the single-cell JAK2/SAT5 model was extended by additional degrees of freedom, which parametrize the covariance matrix of the random effects.

The calibration of the single-cell model was computationally much more demanding, because (i) the additional degrees of freedom increased the dimensionality of the estimation problem, and (ii) the evaluation of the cell-population dynamics required a large number of single-cell simulations, with parameters sampled from the multivariate normal distribution. Hence, we had to improve the feasibility of previously proposed approximation methods. The key challenge was the computational complexity of simulating the population dynamics, a step which needs to be repeated during every step of the model calibration. To reduce the computational costs, the mean and the covariance can be approximated using sigma points (Loos et al, 2018; Toni & Tidor, 2013). However, as shown in Figure 3A, this method possesses a low accuracy for the considered problem. Thus, we introduced a Dirac-mixture distribution (Gilitschenski & Hanebeck, 2013), which allows for a high approximation accuracy via the specification of an appropriate number of samples. The sample points were chosen such that the corresponding Dirac-mixture distribution provided an accurate approximation of the multivariate normal distribution (Hanebeck & Klumpp, 2008) describing cell-to-cell variability (Figure 3B). We used 42 sample points for this approximation, thus we required about 250 times fewer simulations than for a corresponding Monte-Carlo-based trajectory (Figure 3C). Our approach allowed to flexibly balancing the computational complexity, as determined by the number of points, and approximation accuracy. The validity of this approximation, on which all results of the parameter estimation and the model selection relied, was ensured by comparing the final approximate simulations after the model calibration (Figure 3B) to an accurate Monte Carlo simulation with 10 000 samples, which is considered to be as close as possible to the truth (Figure 3C). To further enhance computational efficiency and optimizer convergence, we derived the respective forward sensitivity equations for the gradient calculations, optimized logicle- and log-transformed parameters and used a novel parametrization approach for the covariance matrix (for details, see STAR Methods).

**Figure 3:**
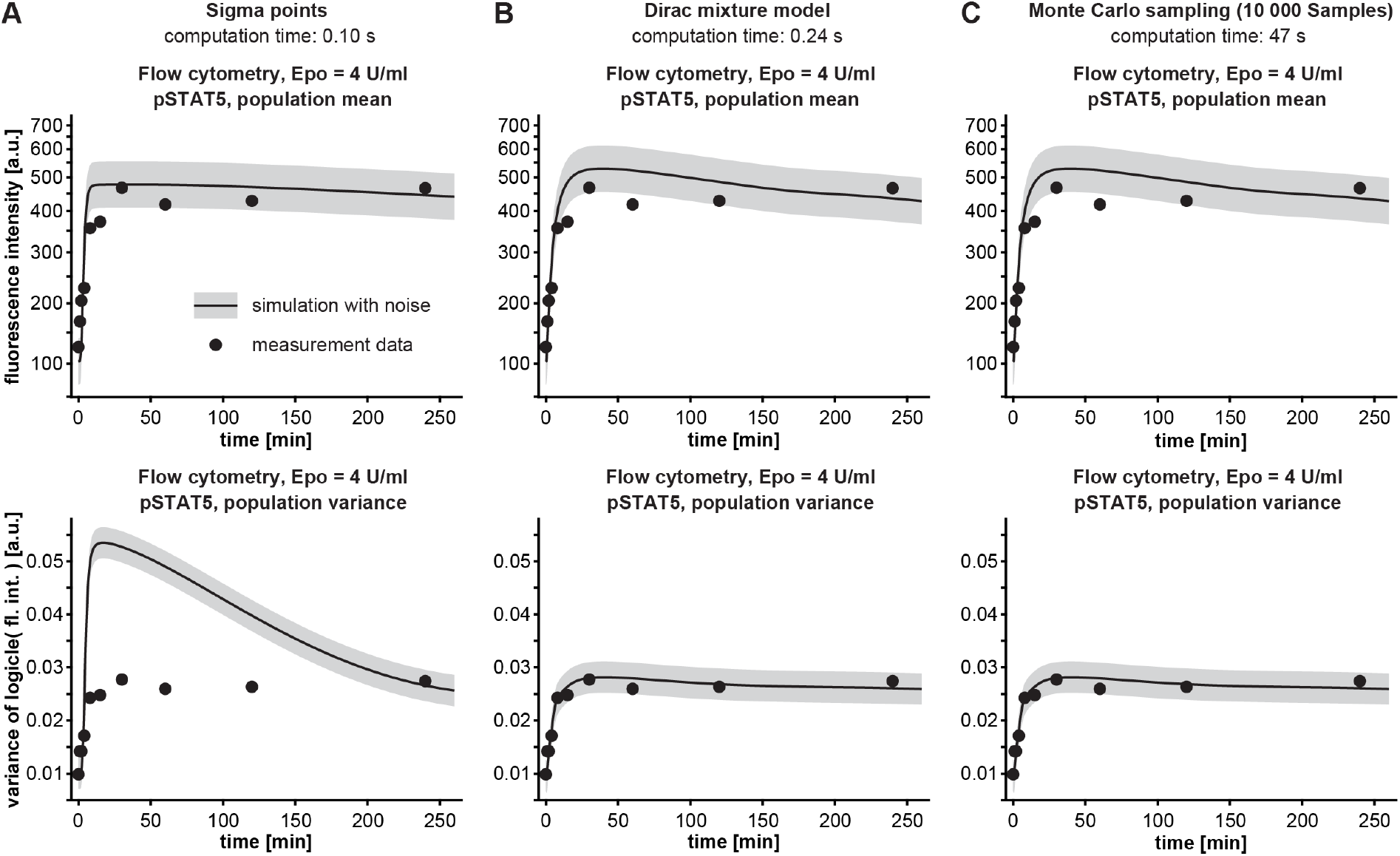
Modeling of population-average dynamics. Exemplary model outputs of different approximation methods for the mean of time-resolved flow cytometry experiment depicted in Figure 2F and the corresponding variance: (A) Sigma points, (B) Dirac-mixture distributions for the chosen number of Dirac points in this study (i.e., 42), and (C) a massive Monte Carlo sampling with 10 000 cells. Means (top panels) and variances (bottom panels) of pSTAT5 for the flow cytometry experiment shown in Figure 2 are plotted for 4 U/ml Epo.

The combination of the improvements reduced computational time by more than 100-fold (Figure 3) compared to available approaches. Nevertheless, parameter estimation using 400 multi-start local optimizations required approximately 10 000 h of computation time for the single-cell model, and therefore parallel computing on large clusters, compared to approximately 85 h for the population-average model. In sum, by developing a novel parameter estimation method for the single-cell model, we enabled its calibration using a combination of population-average and single-cell snapshot data.

### Model selection reveals that abundance of pathway components and nucleocytoplasmic transport rates determine cell-to-cell variability of pSTAT5

Based on the data shown in Figure 1, we hypothesized that the cell-to-cell variability of phosphorylated STAT5 arises from the variability in the abundance of components or reaction rates of the Epo-induced-JAK2/STAT5 pathway that determine the formation of nuclear pSTAT5 in individual cells. Because the high variability of pSTAT5 was already present 15 min after Epo stimulation, when CIS and SOCS3 were not yet expressed (Figure S2 D3/4, E3/4, F5, G3/4, G6), we excluded contributions arising from transcription or translation rates of these negative feedback regulators to the cell-to-cell variability. To identify the relevant components and reactions and to quantify their contribution to the variability of phosphorylated STAT5, we established three nested single-cell models (Single-Cell Model 1-3).

For Single-Cell Model 1 we considered our previously established concept that intracellular information processing is largely determined by the abundance of signaling components (Adlung et al, 2017) and allowed the initial amount of EpoR, SHP1 and total STAT5 to vary between cells. Additionally, the measured basal phosphorylation of STAT5 was represented by a cell-specific offset parameter, which comprised ligand-independent phosphorylation and the background-signal of the secondary antibodies (see Figure 1A). Accordingly, these parameters of an individual cell were a combination of fixed and random effects, while all other parameters were only defined by fixed effects (for details, see STAR Methods). To account for interdependencies between the initial amount of EpoR, SHP1 and total STAT5 as well as the pSTAT5 offset (i.e. basal activation plus measurement background), we estimated variances as well as the full covariance structure of the random effects (Figure 4B, blue).

**Figure 4.**
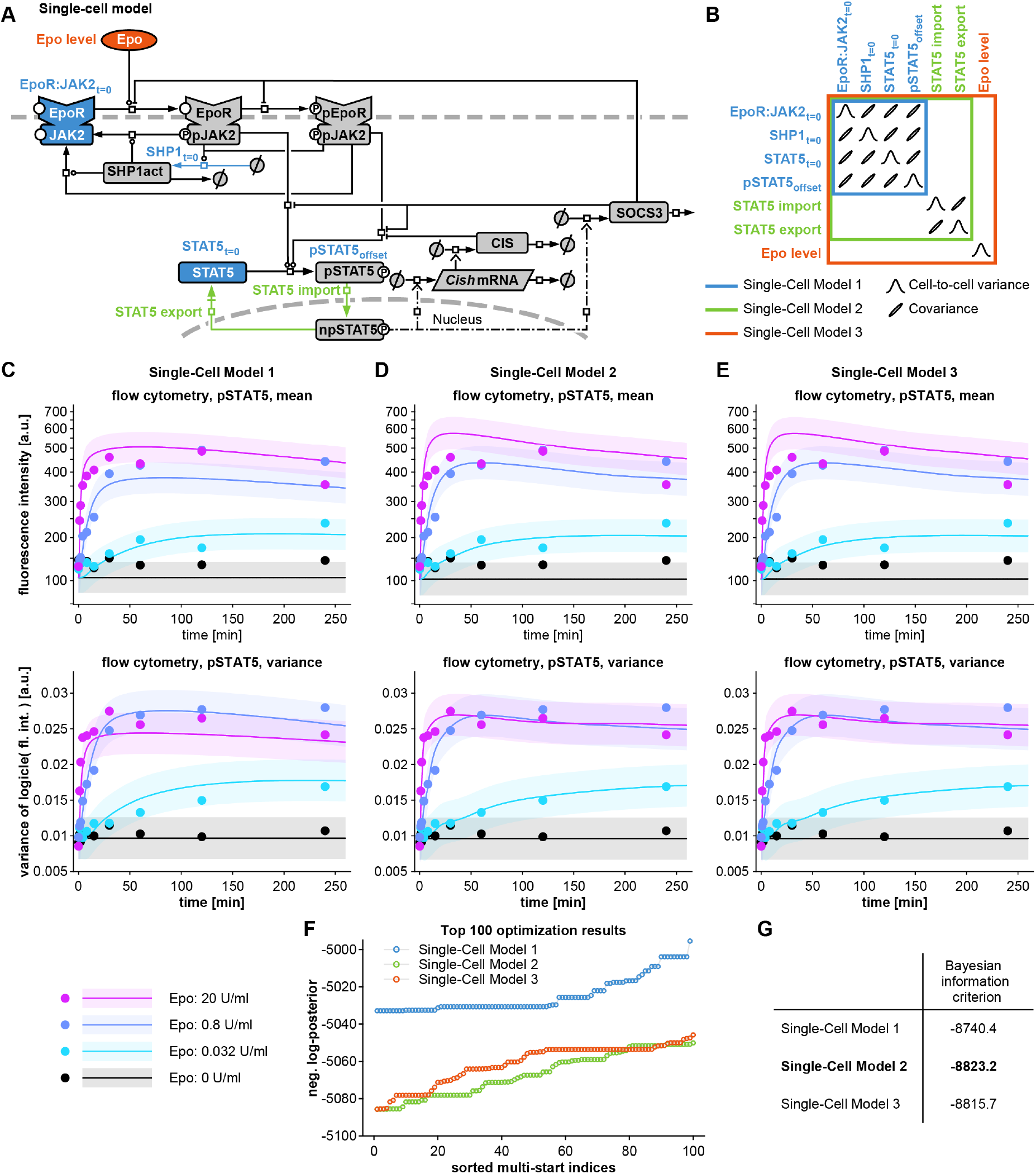
Dynamic mathematical JAK2/STAT5 pathway model describing cell-to-cell variability of phosphorylated STAT5. (A) The structure of the single-cell model of the JAK2/STAT5 pathway with putative sources of cell-to-cell variability of pSTAT5 (colored corresponding to the different models). (B) Visualization of covariance matrix for the three candidate models for the sources of cell-to-cell variability in pSTAT5. (C-E) Experimental data (mean and variance of pSTAT5 from flow cytometry) and mixed-effect model outputs of the three candidate single-cell models. See Figure S4 for correlation of data and model output and Figure S5 for full flow cytometry data set and model output. (F) Multi-start optimization results for candidate single-cell models. The best 100 of 400 local optimization runs are shown, plateaus indicate convergence. (G) Bayesian information criterion (BIC) values of the three candidate models.

In Single-Cell Model 2 we additionally assumed that the nucleocytoplasmic import and export rate constants are variable between cells, since we showed previously that due to the rapid nucleocytoplasmic cycling behavior of STAT5, both the nucleocytoplasmic import and export rates strongly influence the transcriptional yield of nuclear pSTAT5 (Swameye et al, 2003). This could be due to cell-to-cell variability in the cytoplasmic and nuclear volumes of CFU-E cells, which influences the effective import and export rates of individual CFU-E cells (for details, see STAR Methods). We estimated covariances between these two parameters to account for their correlation, but we disregarded possible interdependencies between the translocation rate constants and other entries of the covariance matrix to limit the overall complexity (Figure 4B, green).

In Single-Cell Model 3, we further allowed the input Epo to vary between cells, since we previously observed rapid receptor-mediated ligand internalization (Becker et al, 2010) and therefore considered the possibility of an uneven distribution of the ligand in the medium or differential ligand accessibility of individual CFU-E cells. We considered the random effect of Epo to be independent from the other cell-to-cell variabilities (Figure 4B, orange).

Each of the three single-cell models was calibrated by multi-start local optimization with analytical gradient information of the aforementioned approximation using Dirac-mixture distributions. In addition to the population-average data used for the population-average model (Figure 4C-E, upper panels), the three single-cell models were also parameterized using variances of total STAT5 and pSTAT5 (Figure 4C-E, lower panels), and their covariances, and the means of total STAT5 as measured by flow cytometry, which comprised a total of 960 additional data points. As demonstrated in Figure 4F, the optimizations for all three single-cell models converged well. We performed model selection using the Bayesian information criterion (BIC) (Schwarz, 1978) and found that Single-Cell Model 2 provides an appropriate balance between goodness-of-fit and model complexity (Figure 4G). The optimal values of the log-likelihood functions for Single-Cell Model 2 and 3 were identical, and the difference of 7.5 in the BIC values was due to the penalization of the additional parameters of Single-Cell Model 3. The additional random effect of Epo implemented in Single-Cell Model 3 did not improve the model’s performance. Overall, there was a good agreement and high correlation (*ρ*=0.974, see Figure S4C, D) between model simulations and experimental data (Figure 4C-E), but the single-cell model output was markedly improved by assuming cell-to-cell variability in the import rate of pSTAT5 and the export rate of STAT5 in Model 2 (Figure 4D).

In summary, model selection identified Single-Cell Model 2 that considers cell-to-cell variability for the initial amount of EpoR, SHP1 and total STAT5, as well a cell-specific offset for pSTAT5, and cell specific nucleocytoplasmic import and export rate constants of STAT5, as most suitable to represent the comprehensive experimental data. Thus, we demonstrated that variability in protein abundance of pathway components and in nucleocytoplasmic translocation of STAT5 had a major influence on Epo-induced phosphorylation of STAT5 in individual CFU-E cells and proceeded with this model structure.

### Predictions of cell-to-cell variability in individual cellular states and rate constants

To validate our parameter estimation method based on Dirac-mixture distributions, we compared the single-cell JAK2/STAT5 model output of the distributions of pSTAT5 and total STAT5 to the flow cytometry data for the time-course at 4 U/ml Epo. We observed a good agreement of the mean (Figure 5A, circles) and covariances (Figure 5A, crosses) computed from the flow cytometry measurements and the simulation of the single-cell JAK2/STAT5 model. As these means and covariances were used in the parameter estimation process, this was unsurprising. However, we further compared the population densities, corresponding to two-dimensional histograms with total STAT5 versus pSTAT5 at each time point. Although these densities have not been considered for parameter estimation, the simulated density plots based on the mathematical model and the experimentally determined density plots based on flow cytometry of total STAT5 versus pSTAT5 were comparable (Figure 5A, shades of purple).

**Figure 5.**
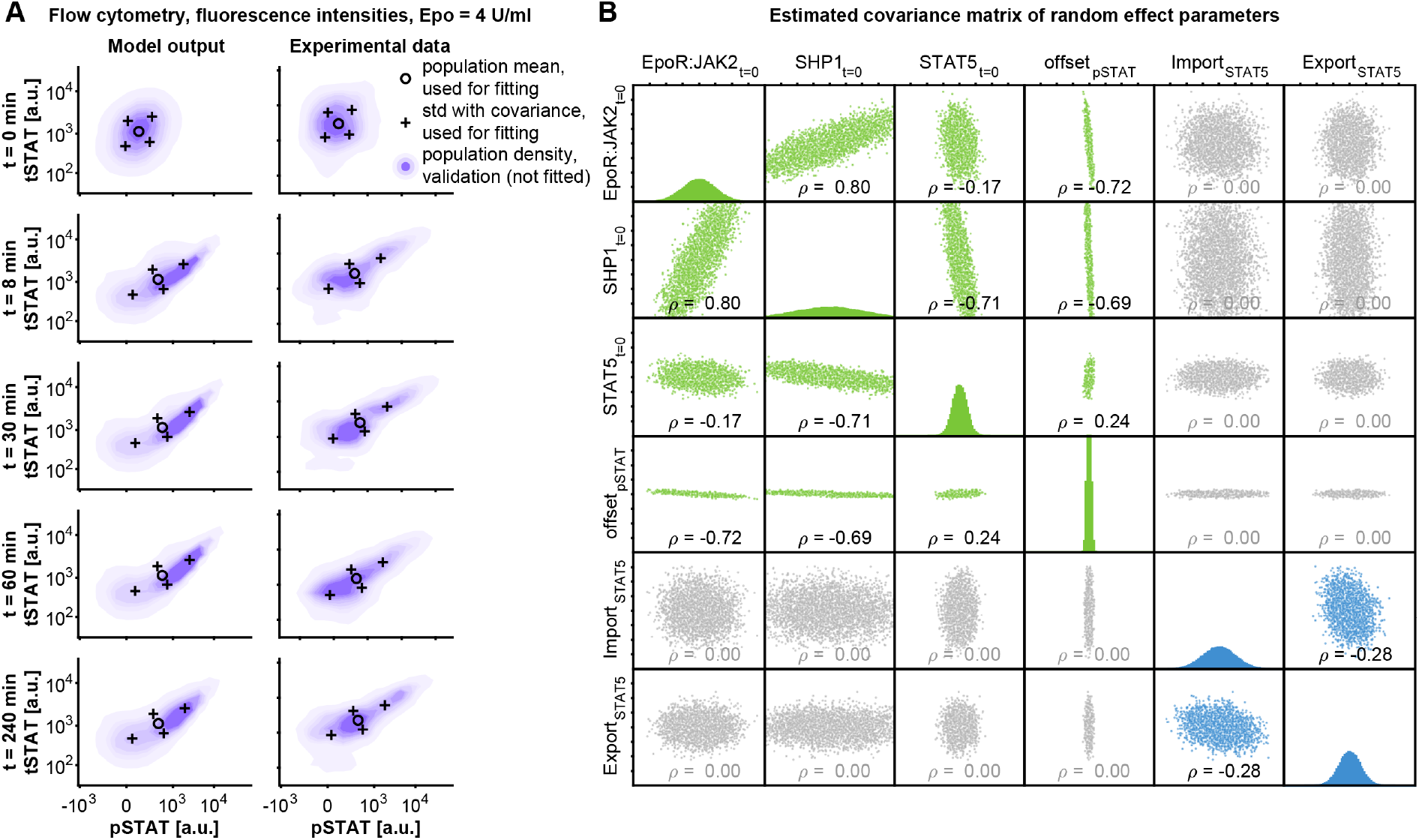
Model simulations of distributions of individual cellular states. (A) Experimental data and output of the single-cell JAK2/STAT5 model for a simulated population of 10 000 CFU-E cells of total STAT5 against pSTAT5. The mean and the covariances of the experimental data shown in Figure S5 that was used for calibration of the single-cell model are shown. Means of model parameters are depicted as circles and the covariances are shown as crosses along the main axes of an ellipse defined by the covariance structure at distances of one standard deviation. The distribution of the pSTAT5 and tSTAT5 intensities of the simulated cell population and the distribution experimentally determined by flow cytometry in a population of 10 000 CFU-E cells are shown as densities. (B) Distributions and corresponding cross-correlations of random effect parameters of the single-cell model.

To identify the pathway components and reactions rates that were responsible for the cell-to-cell variability of phosphorylated STAT5, we plotted the fitted variances and covariances of the random effects (Figure 5B). Based on the model calibration, we predicted higher variances for the membrane-associated EpoR:JAK2 complex and for SHP1 than for total STAT5. Additionally, the inferred covariance matrix revealed correlations between individual cellular states. For instance, the model simulation suggested that the initial abundances of the EpoR:JAK2 complex and SHP1 exhibit a positive correlation of *ρ*=0.8. The positive correlation argued for post-transcriptional co-regulation of the receptor-phosphatase complex for example due to the recruitment of the phosphatase to the phosphorylated EpoR and joint internalization as well as degradation of the SHP1 and the EpoR:JAK2 complex. Further, the model simulations revealed a strong negative correlation between the initial abundance of SHP1 and total STAT5. Because STAT5 is a direct target gene of the JAK2/STAT5 pathway, this negative correlation of protein abundance might be due to transcriptional downregulation of STAT5 in cells with high SHP1 abundance due to enhanced termination of signal transduction through the EpoR.

In addition to correlations of protein abundance, the calibrated single-cell model of the JAK2/STAT5 pathway predicted that the nuclear import and export rates of STAT5 vary substantially between cells. Interestingly, the inferred variance for the nuclear import of pSTAT5 was larger than the inferred variance of the nuclear export of STAT5 (Figure 6A). We quantified this by the ratio of their coefficients of variation, which we assessed by simulating 1000 *in*-*silico* populations with 100 000 CFU-E cells each. This yielded the value

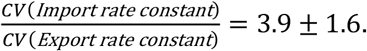

**Figure 6.**
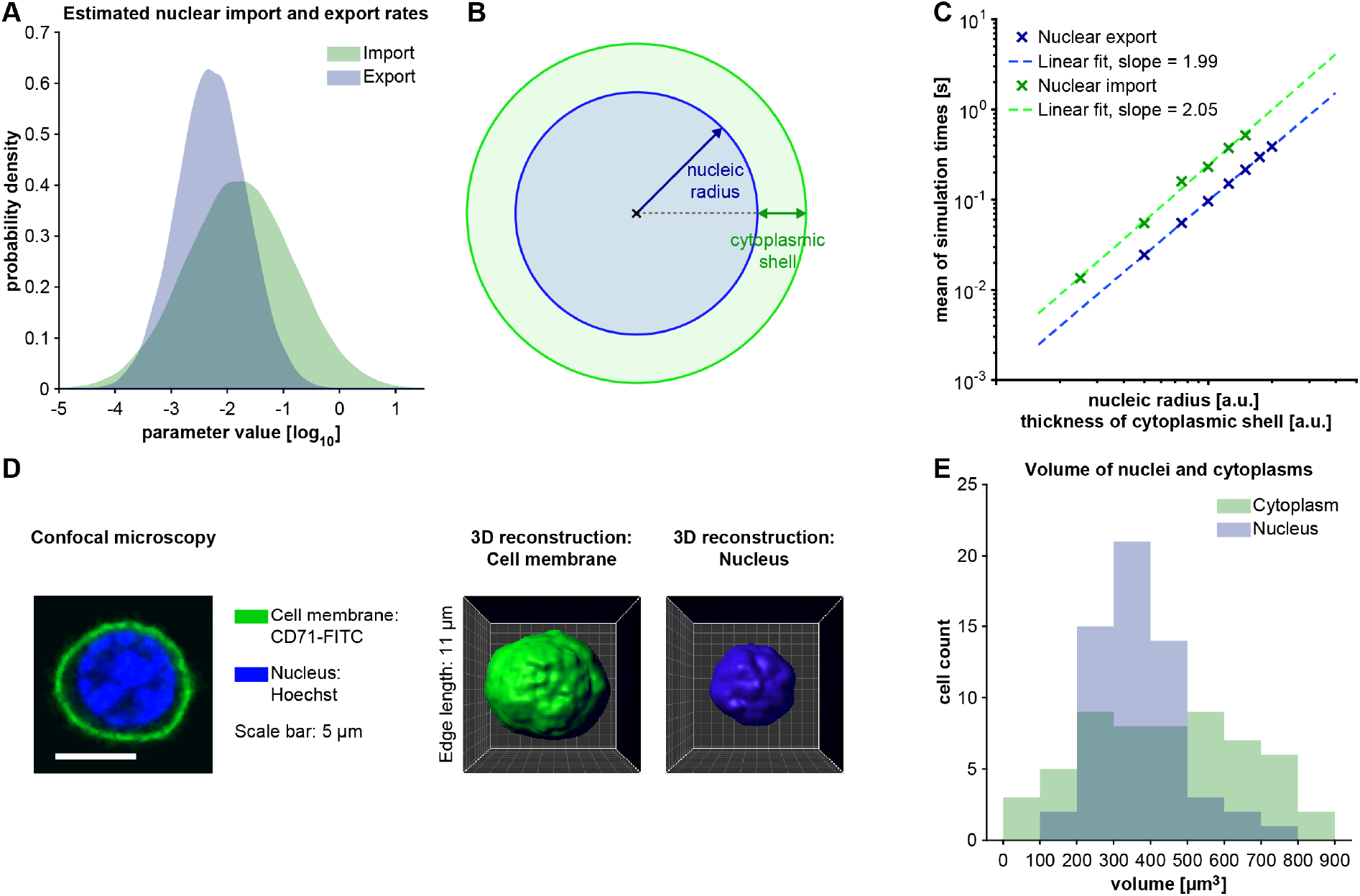
Model simulations of distributions of individual STAT5 import and export rates and experimental validation of the distribution of the nuclear-cytoplasmic translocation of STAT5 in CFU-E cells. (A) Estimated cell-to-cell variability of the import and export rate constants of STAT5 from the single-cell JAK2/STAT5 model. (B) Schematic cross section of a cell indicating the important quantities for nuclear-cytoplasmic translocation rates. (C) Results from a simulation study with 10 000 random walks with fixed step size for different radii of the nucleic sphere core and thicknesses of cytoplasmic shells. (D) The nucleus of unstimulated growth factor-depleted CFU-E cells was stained with Hoechst (Santa Cruz, Catalog #sc-396575), cells were fixed and the cell membrane was labelled with FITC-coupled antibodies against CD71. Z-stack images were acquired by confocal microscopy and used for 3D reconstruction of the cell membrane and the nucleus. (E) Distributions of reconstructed cytoplasmic and nuclear volumes from CFU-E cells are shown (n=58 cells). See Figure S6 for the volume of nucleus and cytoplasm in relation to the volume of the whole cell.

Assuming the transport processes to be driven by diffusion, theoretical considerations (Bressloff, 2014) suggested that the average translocation times of STAT5 (proportional to the inverses of the translocation rate constants) should scale with the squared radius of the nucleic sphere and the squared thickness of the cytoplasmic shell (Figure 6B), respectively (see STAR Methods for more detail). As shown in Figure 6C (see STAR Methods for more details), simulation of these diffusion processes as random-walks in three dimensions and computation of the dependence of expected translocation times of these two quantities (Figure 6C) confirmed this assumptions.

To experimentally validate the prediction of this ratio of variability in the rate constants of STAT5 nuclear translocation, we measured the cytoplasmic and nuclear volumes of CFU-E cells. We stained the plasma membrane and the nucleus of CFU-E cells and reconstructed their 3D structure from confocal fluorescence microscope z-stack images (Figure 6D). In agreement with the predictions of our single-cell JAK2/STAT5 model, the measured nuclear volume of CFU-E cells, which we considered as a proxy for the nuclear export rate constants, showed lower variance than the measured cytoplasmic volume, which we consider as a proxy for the thickness of the cytoplasmic shell and hence for the nuclear import constants (Figure 6E). Furthermore, we quantified the ratio of the coefficients of variation of the corresponding quantities to be

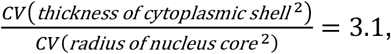

which was in line with our model prediction. In conclusion, we identified that besides STAT5, variability of the EpoR:JAK2 complex and SHP1 as well as of the nuclear import and export rates of STAT5 are responsible for the cell-to-cell variability of STAT5 phosphorylation. The variability of the nuclear import rate is a consequence of the large cell-to-cell variability of the cytoplasmic volumes of CFU-E cells.

### Threshold of nuclear phosphorylated STAT5 determines cell survival

In each individual CFU-E cell survival is ensured by the Epo-induced production of phosphorylated STAT5 that enters the nucleus and induces sufficient amounts of anti-apoptotic gene products. However, it remained unresolved (i) whether the model-predicted cell-to-cell variability of nuclear pSTAT5 was sufficient to explain the experimentally observed survival data, (ii) whether the cells had to acquire a certain amount of nuclear pSTAT5, corresponding to an absolute threshold, or (iii) whether a certain fraction of total STAT5 was required to be phosphorylated and in the nucleus, corresponding to a relative threshold. To address these questions, we established a link between the abundance of nuclear pSTAT5 in each cell given by the single-cell mathematical model of the JAK2/STAT5 pathway to the observed probability for cell survival, depending on the Epo dose. The calibrated single-cell JAK2/STAT5 model was used to simulate the amount of pSTAT5 in the nucleus in each individual cell for different Epo concentrations for time windows between 30 and 180 min. For each of these time windows, the area-under-curve of the nuclear pSTAT5 was calculated for all applied Epo doses and divided by the corresponding time window to obtain the time-averaged abundance of nuclear pSTAT5. Subsequently, the time-averaged abundance of nuclear pSTAT5 was compared to the fraction of surviving cells at the corresponding Epo concentrations. We tested with our single-cell JAK2/STAT5 model if a relative threshold, indicating that a certain fraction of total STAT5 has to be phosphorylated and present in the nucleus to ensure cell survival, or an absolute threshold, meaning that a certain number of nuclear pSTAT5 molecules is required to ensure CFU-E survival, is more consistent with the experimental data. For both single-cell JAK2/STAT5 models we fitted the threshold for different time windows by simulating a population of 10 000 cells and compared the respective goodness-of-fit to the survival data (Figure S7A). The results indicated that the single-cell model realizing a relative threshold to be significantly more informative than the single-cell model with an absolute threshold. Further, the results indicated that the optimal time frame for integrating the amount of nuclear pSTAT5 was the first 120 min after Epo stimulation. The model simulation considering a relative threshold for the nuclear pSTAT5, calculated for a time window of 120 min after Epo stimulation, was in line with the experimentally observed data of the fraction of surviving CFU-E upon stimulation with increasing Epo concentrations (Figure 7A). Additionally, the model simulations predicted that a surprisingly low fraction of about 0.29% of the total STAT5 had to be present as pSTAT5 in the nucleus to ensure the survival of an individual CFU-E cell (Figure 7B and Figure S7B). This means that 24 to 118 pSTAT5 molecules are required in the nucleus of a single cell for 120 min, corresponding to 12 to 59 pSTAT5 dimers.

**Figure 7.**
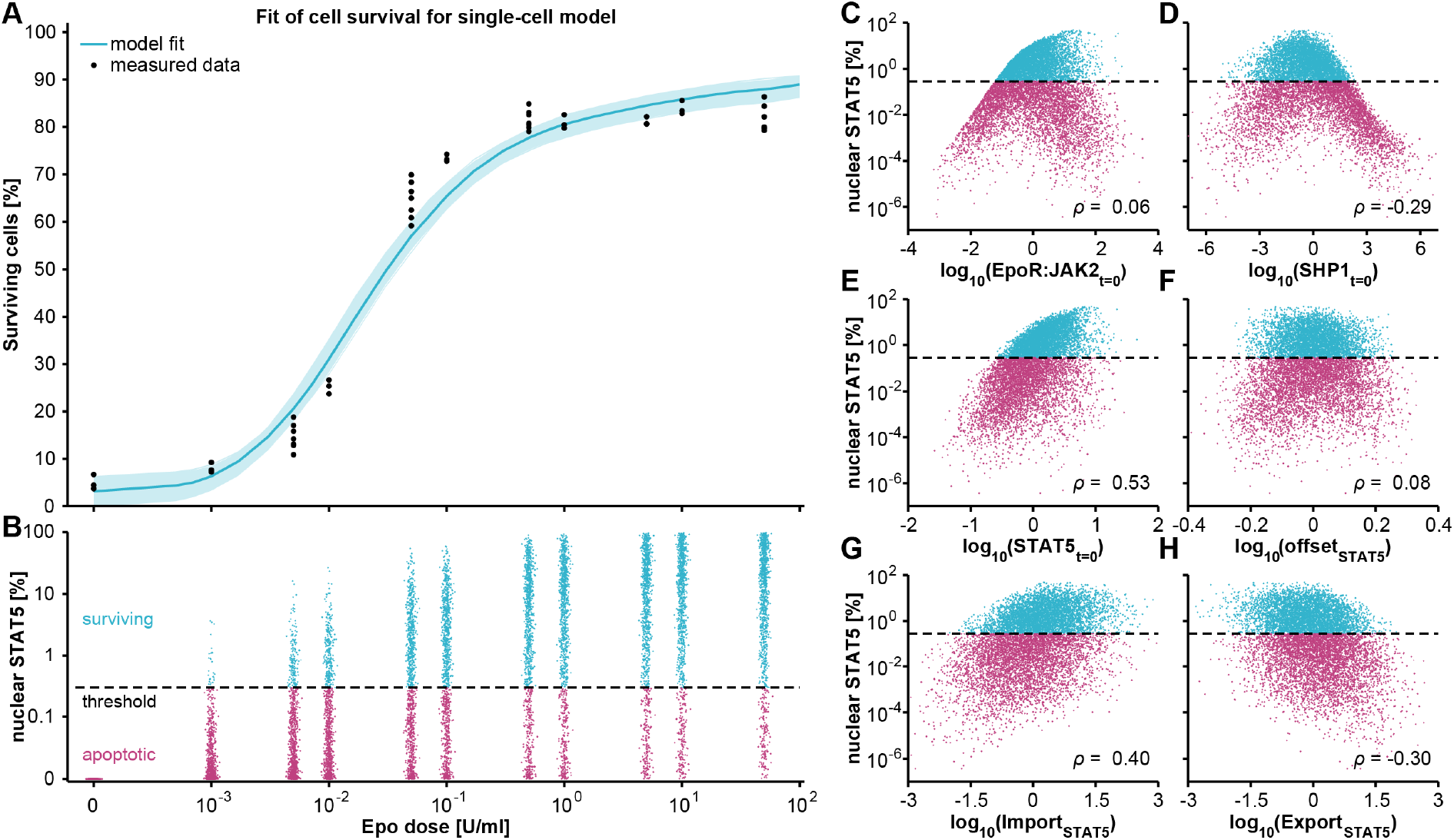
Calibration of the selected JAK2/STAT5 model for cell survival and the influence of random effects on the survival signal. (A, B) Single-cell model calibrated to experimental data of cell survival of CFU-E cells in response to stimulation with the indicated Epo dose. Based on the selected single-cell model, survival criteria based on an absolute or a relative amount of pSTAT5 in the nucleus were compared (see Figure S7). The fit of the survival model based on a relative threshold of 0.29% of total STAT5 during 120 min is depicted as a blue line, the light blue band indicates the model uncertainty. (B) Simulated CFU-E cell populations of the survival model based on a relative threshold of 0.29% of total STAT5 during 120 min, visualized for measured Epo doses. (C -H) Single-cell parameter values for a simulated cell population (Epo at half-effective concentration EC_50_ = 0.032 U/ml) plotted against the fraction of STAT5 in the nucleus of the preferred model. The dashed black line indicates the threshold for cell survival.

To determine which of the components of the JAK2/STAT5 signal transduction pathway with a considerable cell-to-cell variability primarily determine the Epo-dependent life-or-death decision in individual CFU-E cells, the fraction of STAT5 in the nucleus versus the parameter values of these components in single cells was plotted and surviving and apoptotic cells were indicated. While the amount of the EpoR:JAK2 complex (Figure 7C) and SHP1 (Figure 7D) vary substantially in individual cells, only a weak positive correlation of the EpoR:JAK2 complex and a weak negative correlation of SHP1 with the fraction of STAT5 in the nucleus and thus survival was detected. However, for low amounts of EpoR:JAK2, there was a distinct positive correlation with with the fraction of STAT5 in the nucleus, which turned into a negative correlation for higher amounts of EpoR:JAK2. The same effect was observed for SHP1, although here the negative correlation area was more pronounced. This quadratic dependence emerged from a positive correlation of EpoR:JAK2 and a negative correlation of SHP1 with the fraction of STAT5 in the nucleus. A common internalization and degradation of the activated complex and the activated negative regulator might result in this strong correlation of total EpoR:JAK2 and total SHP1 (cf. Figure 5B), which leads to this turning effect. The amount of total STAT5, while not having a large coefficient of variation, positively correlated with the fraction of STAT5 in the nucleus (Figure 7E), while the estimated offset of pSTAT5 in the cytoplasm was not correlated. Interestingly, a positive correlation with the fraction of STAT5 in the nucleus was predicted for the STAT5 import rate (Figure 7G), whereas on the other hand the STAT5 export rate showed a weak negative correlation with the fraction of STAT5 in the nucleus of individual CFU-E cells stimulated with Epo (Figure 7H).

Taken together, we confirmed that cell-to-cell variability in Epo-induced phosphorylated STAT5 is derived from several non-linear regulatory steps. A surprisingly low threshold amount of total STAT5 present as pSTAT5 in the nucleus of an individual CFU-E cell is sufficient to prevent apoptosis and ensure survival of CFU-E cells upon stimulation with Epo. We demonstrated that the STAT5 import rate, which depends on the cytoplasmic volume, had the largest contribution to the fraction of STAT5 in the nucleus and thus to survival of individual CFU-E cells.

## Discussion

In this study, by combining population-level and single-cell data, we discovered multiple sources of cell-to-cell variability in Epo-induced JAK2/STAT5 signal transduction in erythroid progenitor cells. The number of EpoR:JAK2 complexes, as well as the abundance of the phosphatase SHP1 and the cytoplasmic volume are responsible for the variability in STAT5 phosphorylation in CFU-E cells upon Epo stimulation. In addition, we identified a relative threshold of 0.29% of total STAT5 being present as pSTAT5 in the nucleus, which corresponds to as little as 24 to 118 molecules of nuclear pSTAT5 in single CFU-E cells, that is required for the survival of CFU-E cells upon Epo stimulation.

CFU-E cells exhibit the advantage that they rely on Epo as the only factor ensuring cell survival, proliferation and differentiation, which was demonstrated *in vitro* by colony formation (Cooper et al, 1974) and verified *in vivo* using knock-out mice lacking either Epo or its receptor (Wu et al, 1995). Additionally, by deletion of the *STAT5a* and *STAT5b* genes it was shown that survival is primarily controlled by the JAK2/STAT5 pathway (Socolovsky et al, 2001). Binding studies with ^125^I-Epo revealed that murine erythroid progenitor cells exhibit differences in their binding capacities suggesting that the amount of Epo-receptors present on their cell surface is heterogenous (Kelley et al, 1993). We found that variability in the number of EpoR:JAK2 complexes contributes to the cell-to-cell variability in nuclear pSTAT5 in CFU-E cells. In line with our findings, it was shown that variability in the epidermal growth factor (EGF) receptor (EGFR) accounts for cell-to-cell variability in the amount of ERK in the nucleus of rat PC12 pheochromocytoma cells stimulated with EGF (Iwamoto et al, 2016).

Based on theoretical analysis of mathematical models of IFNγ-induced JAK/STAT1 signal transduction it was proposed that the structure of the JAK/STAT pathway renders the pathway relatively insensitive to fluctuations of kinetic parameters and thus ensures robustness (Shudo et al, 2007). Compared to other malignancies of myeloid hematopoiesis, acute erythroleukemia is very rare as it represents only 1% of overall *de novo* AML (Boddu et al, 2018), which could in part be attributed to robustness of the Epo-induced JAK2/STAT5 pathway controlling survival of erythroid progenitor cells. Possibly, because the latent transcription factor STAT5 permits a direct link from the activated EpoR at the cell surface to target gene expression in the nucleus and thereby facilitates cell survival decisions within two hours, the JAK2/STAT5 pathway is less error-prone than more complex pathways with signal amplifications, such as the MAP-kinase pathway that is frequently mutated in human cancer (Dhillon et al, 2007).

While cell fate decisions such as survival or apoptosis are inherently switch-like, at the cell population level the relationship between Epo concentration and fraction of surviving cells shows a graded response curve (Bachmann et al, 2011). Using T cell activation as an example, Zikherman and Au-Yeung recently reviewed how cell-to-cell variability can transform single-cell digital behavior into analog behavior at the population level. At the single-cell level T cells exhibit high variability with respect to their integrated response of T cell receptor signaling. The broad distribution of responses in individual cells leads to a graded dose response at the cell population level and enables the T cell population to distinguish a broad range of antigen dose (Zikherman & Au-Yeung, 2015).

The analysis of our mathematical model identified the hematopoietic tyrosine phosphatase SHP1 as another key component contributing to the observed variability in nuclear pSTAT5 in CFU-E cells upon Epo stimulation. The abundance of SHP1, a phosphatase that is activated by the recruitment to the tyrosin phosphorylated EpoR (Klingmüller et al, 1996), is positively correlated with the amount of the EpoR:JAK2 complex and negatively correlated with the abundance of total STAT5. The positive correlation of the abundance of SHP1 with the amount of the EpoR:JAK2 complex is in line with the mass spectrometric observation that the abundance of proteins forming a common complex is often highly correlated (Wilhelm et al, 2014). The results obtained by metabolic pulse-chase labeling and quantitative mass spectrometry showed that the degradation rates of the free and the bound form differ (McShane et al, 2016) and thus provide a potential explanation for the observed correlation of protein abundance in complexes. Since STAT5 induces its own expression (Yamaji et al, 2013) and the recruitment of the negative regulator SHP1 to the phosphorylated EpoR:JAK2 complex terminates STAT5 signal transduction, heterogeneity in SHP1 levels potentiates the variability in total STAT5 expression across the cell population. This potentiation, together with the heterogeneity in the EpoR:JAK2 complex, contributes to the even higher cell-to-cell variability of pSTAT5 in the nucleus of individual CFU-E cells. Therefore, we speculate that the non-linear impact of SHP1 on both the EpoR:JAK2 complex and on total STAT5 causes that CFU-E cells with an intermediate level of SHP1 are most likely to survive upon Epo stimulation.

Furthermore, we found that the STAT5 import rate, which depends on the cytoplasmic volume, had the largest contribution to the heterogeneity in survival of individual CFU-E cells upon Epo stimulation. In general, nuclear-cytoplasmic translocation rates depend on compartment volumes as proposed by Chara and Brusch (2015). Here, we show a tight coupling between STAT5 transport parameters and cytoplasmic/nucleoplasmic volumes. The large variability in the nuclear import rate of STAT5 in CFU-E cells identified with the model is explained by the experimentally observed large cell-to-cell variability of the cytoplasmic volume of CFU-E cells. This variability in volume might originate from two major sources: It was reported that CFU-E cells can *ex vivo* undergo up to four divisions within 27 h and thus exhibit a rather short cell cycle duration (Nijhof et al, 1984), which might propagate volumetric fluctuations due to rapid successions of cell division. Further, during the differentiation of CFU-E cells to erythroblasts, the cell volume decreases gradually. The proposed interdependency between CFU-E cell cycle progression and differentiation (Pop et al, 2010) could lead to a propagation of the variability that occurs in the cytoplasmic volumes during the G1 phase of the cell cycle, when CFU-E cells have to grow rapidly to reach a critical size for division and differentiation.

The observed larger variability in nuclear import of pSTAT5 compared to nuclear export of pSTAT5 could be explained by recently reported observations suggesting that binding of transcription factors to DNA can act as a passive noise filter and reduce the cell-to-cell variability in nuclear export processes (Battich et al, 2015; Stoeger et al, 2016). The comparison of Epo-induced JAK2/STAT5 signaling between mouse CFU-E cells and the human lung cancer cell line H838 by employing L_1_ regularization to infer cell type-specific parameters in the mathematical model predicted that one of the seven parameters differentially regulated between the cell types was the import rate of pSTAT5 to the nucleus (Merkle et al, 2016) underscoring the potential importance of nuclear import of pSTAT5 for the outcome of the response.

Strikingly, the three components or processes contributing most to the cell-to-cell variability of pSTAT5 in the nucleus are membrane-linked: EpoR:JAK2 and SHP1, which is activated by recruitment to the phosphorylated EpoR (Klingmüller et al, 1996), are localized in the plasma membrane and nuclear import of STAT5 occurs by translocation through the nuclear membrane. Likewise, a previously reported microscopy-based study analysing factors contributing to cell-to-cell variability of simian virus 40 (SV40) infection revealed that the cell-to-cell variability of SV40 infection was caused by differences in the abundance of the sphingolipid GM1 and of the membrane-associated protein focal adhesion kinase (FAK) (Snijder et al, 2009). This indicates that membrane-associated processes frequently exhibit large cell-to-cell variability and are decisive for the regulation of cell fate decisions in single cells.

Our study provides evidence that the threshold of pSTAT5 in the nucleus necessary in the first 120 min after Epo stimulation to ensure survival of a single CFU-E cell is established by a relative fraction of 0.29% of total STAT5. Depending on the cellular amount of total STAT5, this corresponds to 24 to 118 molecules of pSTAT5 present in the nucleus of a CFU-E cell in the first two hours after Epo stimulation. Recently, Frick et al. suggested based on their single-molecule FISH studies of the TGFβ/SMAD signal transduction pathway, in particular the dynamics of SMAD3 nuclear translocation and corresponding gene expression, that relative amounts of transcription factors represent a robust feature to transmit information from the extracellular medium to the nucleus despite cell-to-cell variability in cellular components (Frick et al, 2017). Given our findings this property may also apply to the JAK2/STAT5 signal transduction pathway.

Since we developed a mechanism-based mathematical model of the Epo-stimulated JAK2/STAT5 signal transduction pathway in individual CFU-E cells, the model parameters are directly interpretable. On the contrary, the majority of currently pursued mathematical models remain phenomenological and contain parameters that frequently do not have a direct representation. Data based on single-cell mRNA sequencing in several systems has demonstrated large variability on the mRNA level across cell populations (Buettner et al, 2015; Dalerba et al, 2011). However, it remains unresolved to which extent these differences are translated to variability at the proteome level, in particular since it was reported that the correlation of mRNA expression and protein abundance in mammalian cells can be rather low (Schwanhausser et al, 2011). Furthermore, in these studies dynamic changes in the extent of post-translational modifications are not captured, which are essential for signal transduction and therefore the regulation of cell fate decisions. Currently, to propose temporal ordering and cell fate hierarchies, frequently diffusion pseudotime is estimated from single-cell snapshot expression data (Haghverdi et al, 2016). However, the relationship of this pseudotime to the actual time of each single-cell trajectory is complex. In our study we estimated that a time window of 120 min is required for connecting the nuclear pSTAT5 with cell survival in individual CFU-E cells. This time window estimated for individual CFU-E cells is considerably prolonged compared to the 60 min identified for the cell population to be sufficient to link the integral of pSTAT5 to the survival of CFU-E cells (Bachmann et al, 2011), demonstrating that the time frame of an individual cell can be different from the average population response.

The identification of the causes and consequences of cell-to-cell variability in the JAK2/STAT5 signal transduction pathway in CFU-E cells was only possible by developing a mechanism-based single-cell model that required the combination of experimental data obtained at the population-average and at the single-cell level together with the identification of Dirac-mixture distributions as a suitable approximation approach. Similar to signal transduction addressed in this study, transcriptional regulation can now be experimentally addressed using genome-wide population-average measurements and single-molecule techniques and also there it was realized that it is crucial to combine these information levels and consider the binding energy of transcription factors to DNA as well as the different rates of transcription initiation to develop a quantitative model of transcriptional regulation (Coulon et al, 2013).

In conclusion, we present a two-step mathematical modeling strategy to integrate population average data and single-cell data to determine a threshold for binary cell fate decisions. This approach is scalable and can readily be applied to other cell fate decisions such as proliferation, migration and differentiation that are controlled by multiple transcription factors in health and disease.

## STAR Methods

Detailed methods are provided in the online version of this paper and include the following:

- Key resource table
- Contact for reagent and resource sharing
- Experimental model and subject

- Preparation of erythroid progenitor cells
- Confocal microscopy and data analysis
- Flow cytometry
- Cell fractionation
- Quantitative immunoblotting, qPCR and mass spectrometry
- Methods Details

- Mathematical modeling of population-average data
- Mixed-effect modeling for cell population
- Approximations to simulate cell populations
- Parametrizations of covariance matrices
- Parameter estimation for single-cell model
- Quantification and statistical analysis

- Model selection criteria
- Translocation rates and compartmental volumes
- Inference of the survival criterion
- Data and software availability

## Supplemental information

Download link to supplementary file

## Acknowledgments

C.T. and J.T. acknowledge support by the state of Baden-Württemberg, Germany, through bwHPC and the German Research Foundation (DFG) through grant INST35/1134-1 FUGG. P.S. and J.H. acknowledge support by the European Union’s Horizon 2020 research and innovation program (CanPathPro; Grant No. 686282). L.A., C.T., J.H., U.K. and M.S. acknowledge funding by the German Ministry of Education and Research (BMBF) within the e:Bio collaborative research projects “Systems Biology of Erythropoietin” (SBEpo, 0316182A and 0316182B). J.T., M.S. and U.K. were supported by the German Federal Ministry of Education and Research (BMBF)-funded ERAPerMed consortium “Improved Treatments of Acute Myeloid Leukaemias by Personalised Medicine” (AML_PM, 01KU1902A and 01KU1902B) J.T. and U.K. were supported by the German Federal Ministry of Education and Research (BMBF)-funded e:Bio consortium MS_DILI (research grants 031L0074B and 031L0074A) and the Systems Medicine network LiSyM (research grants 031L0048 and 031L0042). C.T and J.T. were supported by the Deutsche Forschungsgemeinschaft (DFG, German Research Foundation) under Germany’s Excellence Strategy -EXC-2189 -Project ID: 390939984.

## Author contribution

M.S., J.H., U.K. and J.T. designed the study. L.A. conducted and analyzed all experiments except for the microscopy conducted and analyzed by L.E.S.. C.T. and J.T. implemented, calibrated and reduced complexity of the ODE model of the population average data. P.S. carried out the model selection for the single-cell models and developed and implemented the parametrization method for the covariance matrices. P.S., L.S. and J.H. implemented and calibrated the single-cell model. P.S., D.W., and J.H. implemented the Dirac mixture distribution. L.A., C.T., P.S., L.S., J.H., U.K. and M.S. wrote the manuscript, with input from J.T. All authors read and approved the manuscript.

## STAR Methods

### KEY RESOURCE TABLE

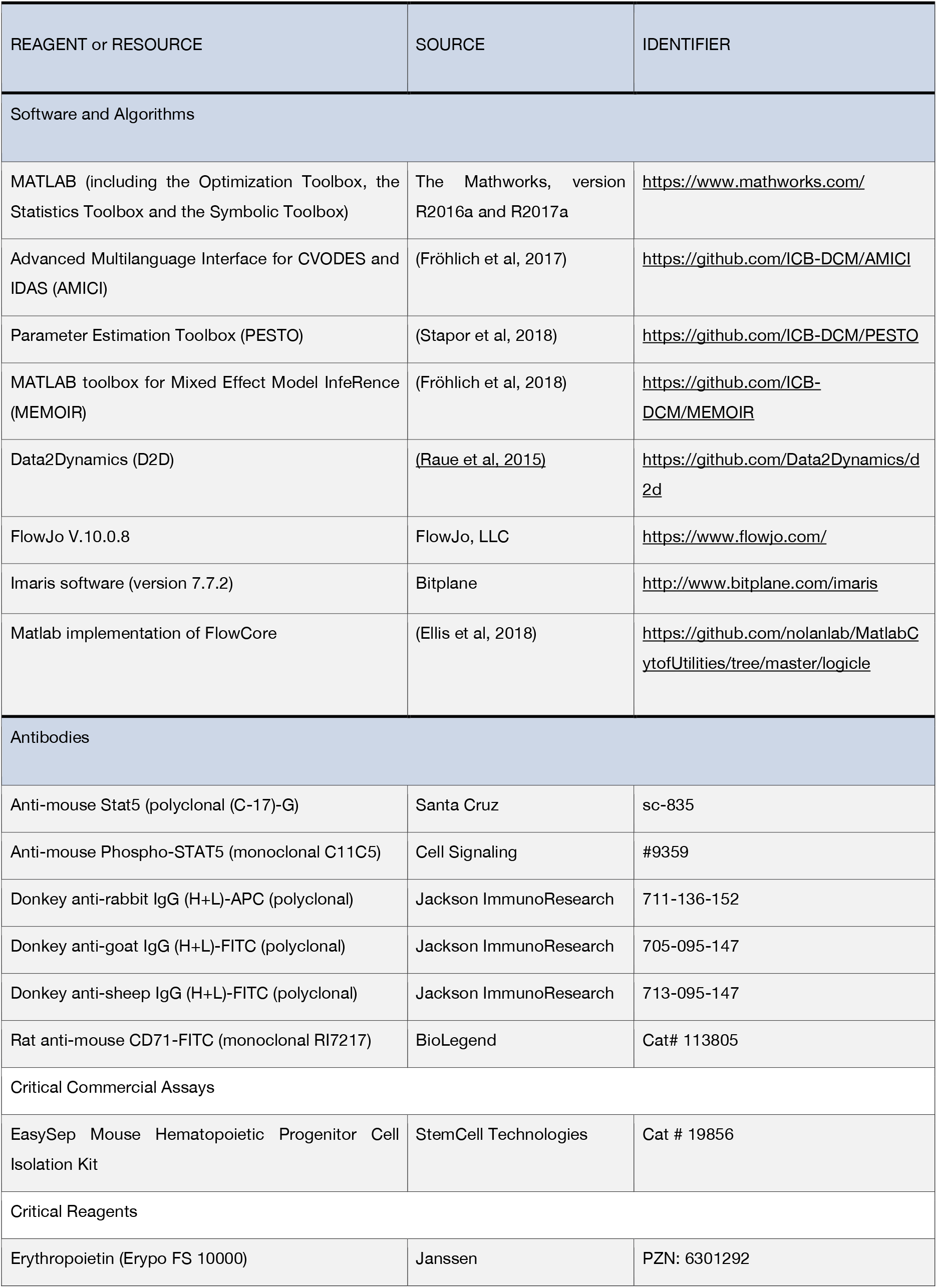
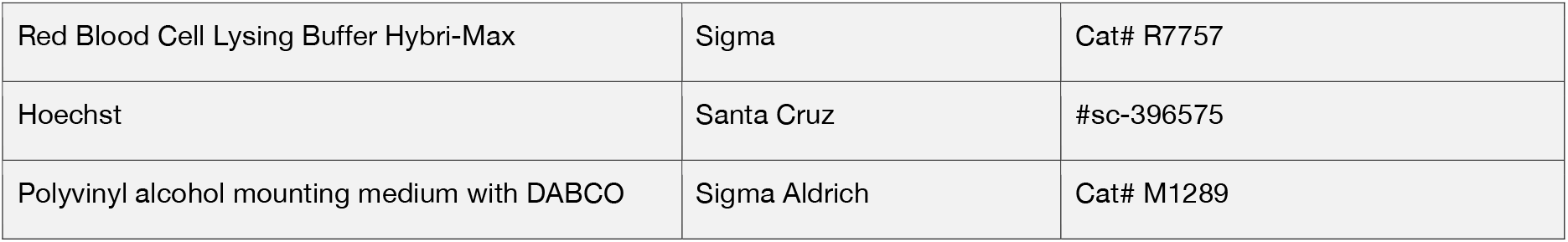

### CONTACT FOR REAGENT AND RESOURCE SHARING

Further information and requests should be directed to the Lead Contact Marcel Schilling.

### EXPERIMENTAL MODEL AND SUBJECT

#### Preparation of erythroid progenitor cells

All mice of this work were housed at the DKFZ animal facility under constant light/dark cycles. Animals were maintained on a standard mouse diet and allowed *ad libitum* access to food and water. All animal experiments were approved by the governmental review committee on animal care of the federal state of Baden-Württemberg, Germany (reference number DKFZ215).

Fetal mouse livers at embryonic day 13.5 (i.e. during massive erythroid expansion) were isolated from the uteri of sacrificed Balb/c mice (Potter, 1985). Fetal liver cells (FLCs) were resuspended in 500 μl of 0.3 % BSA/PBS, flushed through a 40 μm cell strainer (BD Biosciences) and taken up in 10 ml Red Blood Cell Lysing Buffer (Sigma-Aldrich). The suspension of 40 fetal livers was subjected to negative depletion by using magnetic beads (Miltenyi Biotech). Freshly purified CFU-E cells were cultivated for 14 h in Panserin 401 (PAN-Biotech) supplemented with 50 μM 2-mercaptoethanol and 0.5 U/ml Epo. Subsequently, CFU-E cells were washed three times with Panserin 401 supplemented with 50 μM 2-mercaptoethanol and deprived from growth factors in Panserin 401 supplemented with 50 μM 2-mercaptoethanol and 1 mg/ml BSA at 37°C for 1 h.

#### Confocal microscopy and data analysis

Staining of the nucleus was performed with 2 μg/ml Hoechst (Santa Cruz, Catalog #sc-396575) at a cell density of 10×10^6^ cells/ml for 20 min at 37°C, shaking at 800 rpm. The cells were fixated in 4% formaldehyde, followed by staining of the plasma membrane with 15 μg/ml FITC-coupled rat anti-mouse CD71 (Biolegend, Cat# 113805) for 30 min at room temperature. For microscopy the cells were mixed 1:1 with Polyvinyl alcohol mounting medium with DABCO (Sigma Aldrich, Cat# M1289).

Confocal microscopy was performed with the ZEISS LSM 710 ConfoCor 3 using a 43x magnification oil objective. FITC and Hoechst (Santa Cruz, Catalog #sc-396575) were excited by a 488 nm and 405 nm Argon Laser, respectively. Z-stacks with an interval of 0.5 μm were taken and subsequently three-dimensionally reconstructed with the ImageJ software (version 1.4.3.67). The nuclear and cytoplasmic volumes were calculated by artificially filling the volume enclosed by the stain using the Fill Holes and 3D Objects Counter commands. For illustrative purposes, the cells were three-dimensionally reconstructed with the Imaris software (version 7.7.2) and the shape of the plasma membrane and the nucleus were highlighted with the Surface function.

#### Flow cytometry

Phosphorylated and total STAT5 was stained as follows: Stimulated CFU-E cells were fixed in Fixation Buffer (BD Biosciences) or 4% PFA in PBS. Cells were permeabilized in ice-cold Perm Buffer III (BD Biosciences) or 90% methanol. Prior to acquisition, cells were washed with Stain Buffer (BD Biosciences) or 0.3 % BSA/PBS. pSTAT5 was detected with rabbit anti-mouse phospho-STAT5 antibodies, total STAT5 was detected with goat anti-mouse Stat5 antibodies. The secondary antibodies were anti-rabbit coupled to APC for pSTAT5, and anti-goat coupled to FITC for total STAT5.

Flow cytometry was performed on a LSR II flow cytometer (BD Biosciences). Recorded files were exported and subsequently analyzed with FlowJo V.10.0.8.

#### Cell fractionation

For fractionation of cytoplasmic and nuclear lysates, CFU-E cells were treated with 2× Homogenization buffer 1 (1× buffer: 0.1mM EDTA, 0.1mM EGTA, 10mM NaF, 10 mM HEPES pH 7.9, 10 mM KCl, 1 mM DDT, 1 mM NaV_3_VO_4_) supplemented with 2 μg/ml aprotinin and 200 μg/ml AEBSF. Upon addition of 1× Homogenization buffer 1 supplemented with 5 % NP-40, samples were incubated for 5 min on ice and then centrifuged for 3 min at 14,000 rpm and 4°C. Supernatant was taken as cytoplasmic fraction. Remaining pellets were washed three times with 1× Homogenization buffer 1 and subsequently re-suspended in 1× Homogenization buffer 2 (400 mM NaCl, 1 mM EDTA, 1 mM EGTA, 10 mM NaF, 10 mM HEPES pH 7.9, 1 mM DDT, 1 mM NaV_3_VO_4_) supplemented with 1 μg/ml aprotinin and 100 μg/ml AEBSF. Samples were shaken at 1,400 rpm and 4°C for 1 h, sonicated for 30 seconds and then centrifuged for 20 min at 14,000 rpm and 4°C. The supernatant was taken as nuclear fraction.

#### Quantitative immunoblotting, qPCR and mass spectrometry

Quantitative immunoblotting, qPCR and mass spectrometry was performed as described earlier (Bachmann et al, 2011).

## METHOD DETAILS

### Mathematical modeling of population-average data

#### Model topology and data integration

The published ordinary differential equations (ODEs) and the corresponding experimental data (Bachmann et al, 2011) were used as a basis for the mathematical modeling. To ensure comparability on absolute scales with the mixed-effect models, parameter transformations implemented in the published model were avoided. To focus the analysis, we neglected the nine data points for the CIS overexpression condition and removed the mechanism of unspecific binding of CIS to EpoR in the absence of Epo. These adaptions reduced the computational effort when simulating the model output without dropping any other key features of the original model. Complementary, the error of relative pSTAT measured by mass spectrometry was kept constant to 5.55% as previously defined (Boehm et al, 2014; Hahn et al, 2013).

The fluorescence intensities recorded in the flow cytometry experiments were transformed to logicle scale, which is a biexponential scale (Herzenberg et al, 2006; Moore & Parks, 2012; Parks et al, 2006):

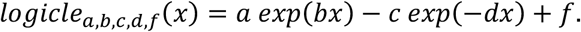

The coefficients *a*, *b*, *c*, *d*, and *f* are determined from the following quantities, which only depend on the data set:

- *T*: Top of the data scale. We used *T* = 2^18^.
- *W*: Width of the data range in approximately linear scale in decades. We used *W* = 0.74 as suggested in Moore and Parks (2012).
- *M*: Width of data range in approximately logarithmic scale in decades. We used *M* = 4.5, as suggested in Moore and Parks (2012).
- *A*: Additional width of data range in negative scale. We used *A* = 0, which reproduces the standard logicle scale as introduced in Parks et al. (2006).

We employed a MATLAB implementation which determines the coefficients *a*, *b*, *c*, *d*, and *f* directly from *T*, *M*, *W*, and *A* according to Moore and Parks (2012).

Utilization of this scale is common practice for the visualization and statistical analysis of flow cytometry data and is specifically useful for cases in which negative values appear after background correction of the raw data and when a more accurate representation of the distribution of intensities around zero is favorable. While the traditional logarithmic scaling of the fluorescence intensities yields a clear data distribution for higher values, it cannot cope with negative values and it tends to ‘piling up’ of intensities on the lower end of the scale. By construction, the logicle scale is able to combine the advantages of the logarithmic scale for higher values with a linear-like scale for the lower end.

For each time point and Epo dose the mean of the logicle scale-transformed fluorescence intensities was used as data point for the fitting of the population-average model. For the quantitative immunoblotting, real-time PCR and mass spectrometry experiments the measured intensities, which should correspond to the mean of the measured cell population, were considered as data points.

#### Parameter estimation and profile likelihood

All calculations and analyses of the ODE system describing the pathway model were done within the MATLAB-based modeling environment D2D (www.data2dynamics.org) (Raue et al, 2015). The ODEs were solved by a parallelized implementation of the CVODES initial value problem solver (Hindmarsh et al, 2005). The ODE model was calibrated using maximum likelihood estimation (Raue et al, 2015). We used the local deterministic Gauss-Newton gradient-based trust-region optimizer implemented in the MATLAB function lsqnonlin which is part of the MATLAB optimization toolbox. Multi-start optimization runs with in each case at least 2500 initial guesses were performed using the afore-described model and dataset as well as for each reduced model (see section “Model reduction”). To assess the uncertainty and identifiability of the estimated parameters the profile likelihood (Raue et al, 2010; Raue et al, 2009) was calculated using D2D.

#### Model reduction

Based on the comprehensive experimental data set and the original population-average model topology, a systematic data-based model reduction was performed by iteratively analyzing the profile likelihood (Hass et al, 2017; Maiwald et al, 2016; Tönsing et al, 2018). By this, non-identifiabilities can be resolved without changing the dynamics of the observed model entities.

First, we observed the inverse coupling of the non-identifiable parameters for the activation rate of SHP1 by the active form of the EpoRJAK2 (EpoRpJAK2 and pEpoRpJAK2) complex and the rate by which the pEpoRpJAK2 state is recycled to its deactivated form by activated SHP1 (SHP1Act). This suggests a weak activation of SHP1. Analogously to scenario 3 in Maiwald et al. (2016), the SHP1 activation is so weak that it merely changes the level of the inactive SHP1 state, although the amount of active SHP1 is not negligible, as it mediates the deactivation of the EpoRJAK2 receptor. As a result, the profile likelihood of the activation rate is open to minus infinity on the log_10_ scale, i.e. to zero on the linear scale, while the SHP1Act mediated deactivation rate compensates for this and its profile is open to infinity. We resolved this non-identifiability by reparametrizing the SHP1 mediated deactivation rate with the inverse of the SHP1 activation rate, which yields the new, practically identifiable parameter DeaEpoRJAKActSHP1 and structural non-identifiability of the SHP1 activation rate. The latter is resolved by fixing it to an arbitrary value, which in the end is equivalent to a constant SHP1 state with fixed functional relation between SHP1 and its active form.

Secondly, we observed the non-identifiability of the CISmRNA turnover parameter (CISRNATurn) with a profile likelihood open to infinity. This parameter is related to the modeling of a transcriptional delay in the synthesis of CISmRNA via the linear chain trick (MacDonald, 1976). We resolved this non-identifiability by iteratively reducing the chain length from five to two.

Thridly, we observed that the SOCS3mRNA transcription process showed a similar behavior in the SOCS3mRNA turnover parameter (SOCS3TurnRNA) and the corresponding delay parameter (SOCS3RNADelay), as well as the SOCS3 translational process in the parameter (SOCS3Turn). All three model parameters showed a practically non-identifiability with profile likelihoods open to infinity. This implies that the measured data of SOCS3 mRNA and SOCS3 protein can be described by the model without delay but a single step. This is consistent with the experimental data for SOCS3 mRNA and SOCS3 protein, which do not indicate a delay. As a consequence, the transcriptional delay, the transcriptional and the translational process of the synthesis of SOCS3 were merged into one single reaction.

The three model reduction steps yielded a model with only two remaining practically non-identifiable parameters: the rate of Epo receptor mediated STAT5 phosphorylation (STAT5ActEpoR) and the parameter capturing the inhibition STAT5 phosphorylation by CIS (CISInh). These parameters cannot be determined from the available experimental data, but related studies showed the relevance of these processes (Gobert et al, 1996; Yoshimura et al, 1995). To ensure biological plausibility of the model and because these non-identifiabilities do not diminish the predictive power of the model's output, we did not perform an additional model reduction step.

The different parameter and state eliminations provide a reduced model (see model equations below) which captures the core features of the original model and is able to describe all available population-average data. However, the identifiability of the remaining model parameters improved considerably when compared to the model suggested from Bachmann et al. (2011) and provides finite profile likelihood based confidence intervals at a confidence level of 0.95 for 19 of the 21 dynamic parameters. At a confidence level of 0.68, all dynamic parameters have finite profile likelihood-based confidence intervals. In addition, the described model reduction resulted in a substantial improvement of optimizer performance and convergence. On average 4.1 fits per hour converged to the global optimum for the reduced model and the complete data set, whereas only 2.6 fits per hour converged to the global optimum for the initial model structure. All fits belonged to a multi-start sequence with 5000 initial guesses on a 16-core @ 2.4 GHz CPU.

#### Model equations

Set of ordinary differential equations (ODEs) for the reduced model:

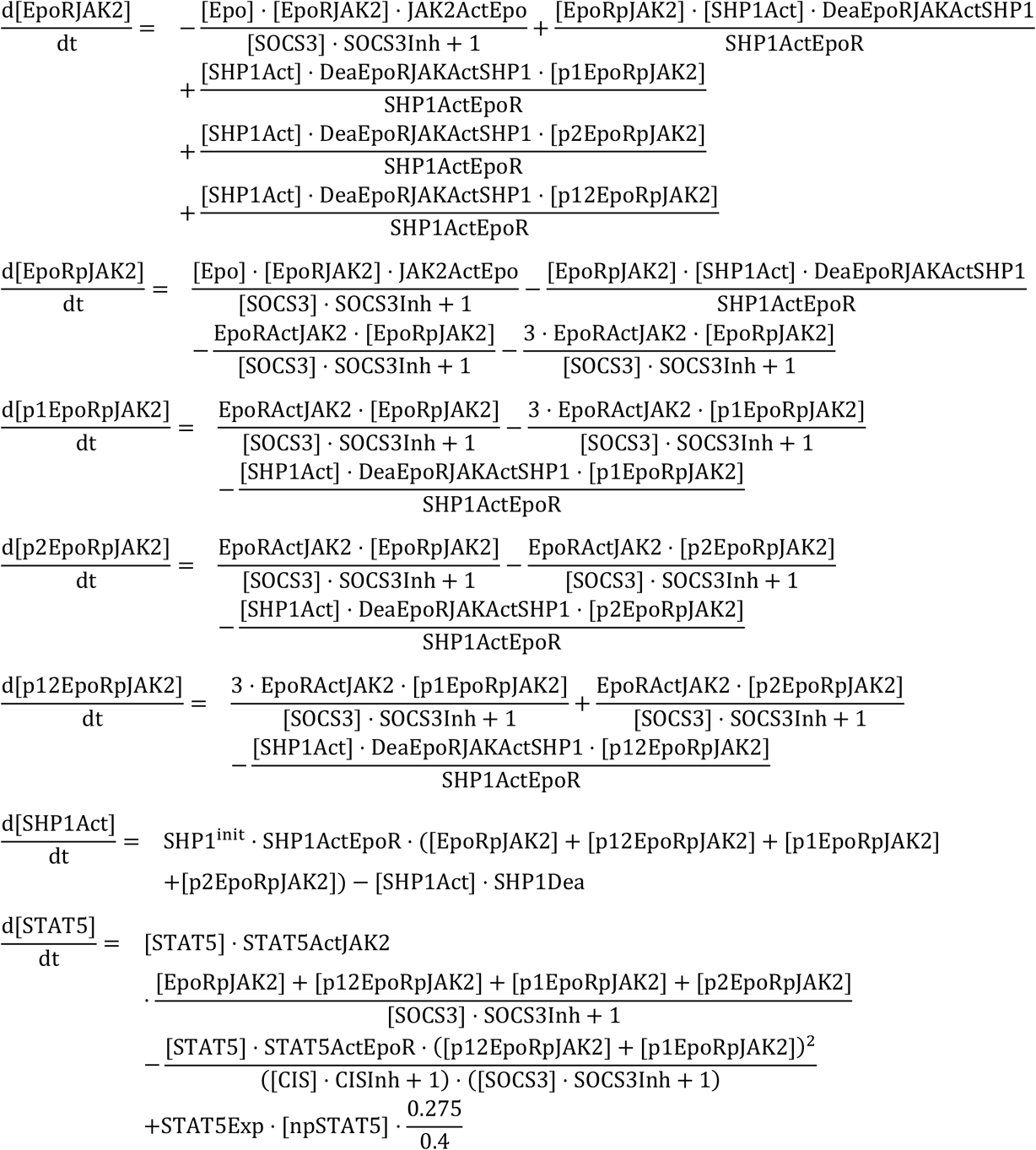

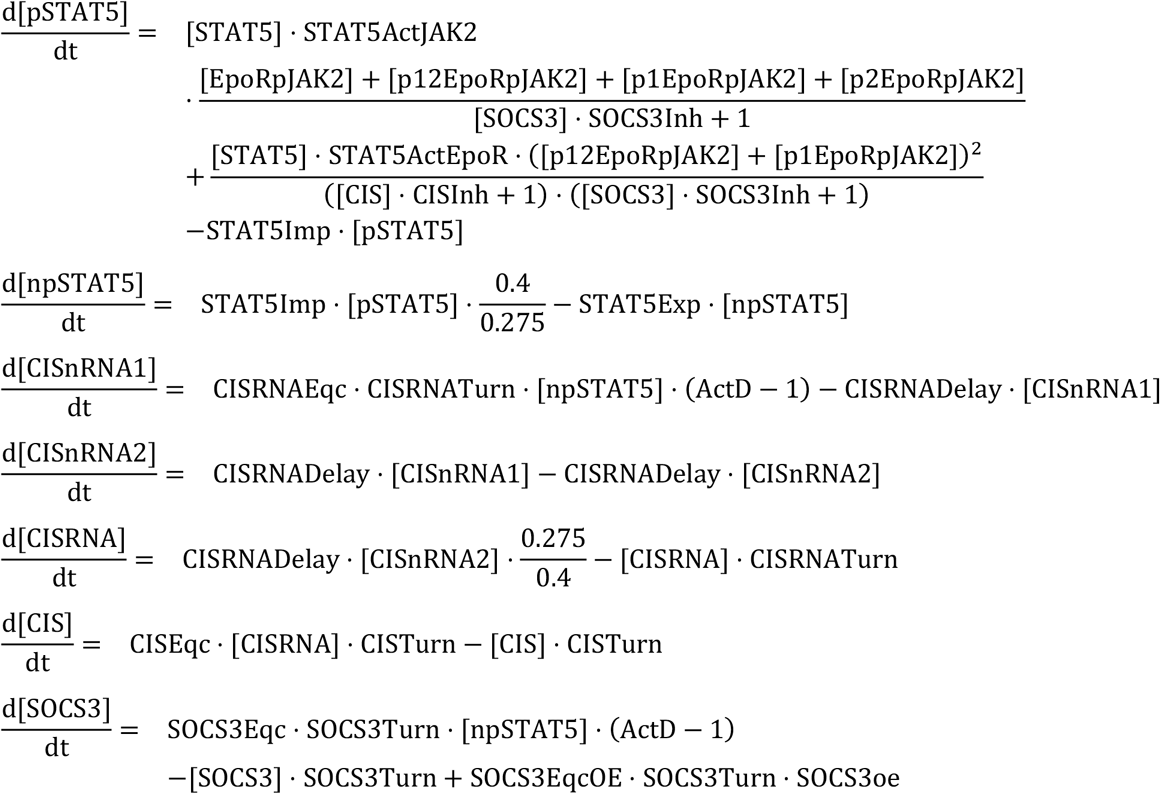

Initial values of ODE system:

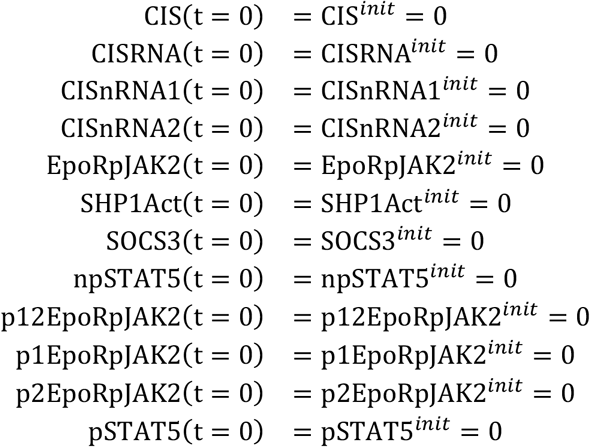

Applied parameter transformations:

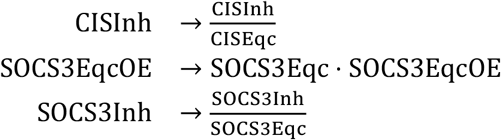

### Mixed-effect modeling for cell population

In this study we account for cell-to-cell variability be allowing the protein abundances and parameters of individual cells to differ. This is a common approach to capture (slow) extrinsic noise. As the abundance of most biochemical species involved in the process is rather high, we assume that the influence of intrinsic noise (stochasticity of cellular processes causes differences) is minimal. Mathematically, considering differences between the parameters of individual cells yield a mixed-effect model of the process. As we propagate this model through an ODE - providing a non-linear map from parameters to outputs -, we obtain a nonlinear mixed-effect model (NLMEM).

To simulate the NLMEM, we create an *in-silico* population of cells, in which each single-cell has its own parameter vector. The parameter vector *ϕ*^*i*^ for the i-th cell is given as

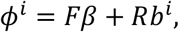

in which *F* and *R* are called design matrices for fixed and random effects, 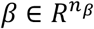 is the vector of fixed effects (i.e., the parameter set describing the population mean),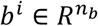 is the vector of random effects for the i-th cell. The fixed effects *β* influence the parameters of all cells, while the random effects *b*^*i*^ are specific to one single-cell. We assume the random effects to follow a multivariate normal distribution with mean 0 and covariance matrix, *Σ*, *b*^*i*^ ~ *N*(0, *Σ*). The covariance matrix *∑* is parameterized by a vector *δ*, yielding the parameter vector *θ* = (*β*, *δ*), which describes the population dynamics. Hence, the parameter estimation problem of the population average model (where only *β* has to be inferred) is extended for the mixed effect model by the parameters of the population distribution which are grouped in *δ*. The term “NLMEM” is typically used when analyzing single-cell time-lapse data (Karlsson et al, 2015), however, many of the models used for single-cell snapshot data are also NLMEM (Loos et al, 2018). As the cells from different time points in a time series of flow cytometry data are not the same and are hence not directly comparable, we did not fit single-cell trajectories to the whole dataset. Instead, we simulated a heterogeneous population, propagated it through the ODE and fitted the shapes of the simulated distributions of certain protein abundances to the snapshots. As a measure of the shape of these distributions, we used means, variances and covariances.

In more detail, this means that for a population with *M* individuals, we simulated *M* trajectories of state variables

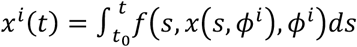

for each random effect vector *b*^*i*^, where *f* is the vector field of the considered ODE model: *x*^*i*^(*t*) = *f*(*t*,*x*^*i*^(*t*),*ϕ*^*i*^) with *x*^*i*^(*t_0_*) = *x_0_*(*ϕ*^*i*^). This yielded *M* trajectories of observables

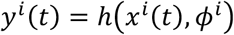

from which the means, variances and the covariances (and possibly higher moments) can be computed and fitted to measurement data.

### Approximations to simulate cell populations

#### Monte Carlo sampling

The simplest way to approximate the means, variances and the covariances is to directly simulate a cell population using parameters obtained from Monte Carlo sampling. For each individual cell ***i*** ∈ **1**, …, ***M***, the random effect vector ***b***^*i*^ is given by

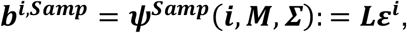

in which ***L*** is the lower factor of the Cholesky decomposition of covariance matrix ***Σ*** and ***ε***^*i*^ is th *i*-th sample from a set of ***M*** standard normally distributed points in 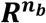. Outputs like mean and covariance of observables are obtained from the outputs of each sample.

The size ***M*** of the sampled population must be large enough to reflect the behavior of the distribution function. For this purpose, sample sizes of e.g. 10 000 simulated cells are common, which means that the computation time necessary for model parametrization must be multiplied with the size of the cell population. While this may be feasible for analytic or very simple ODE models, it is computationally intractable for the model considered in this work, since this would result in millions of hours of computation time. However, simulating a full population of cells represents the most faithful method for describing the actual biological system. Hence, we compared the final simulation results obtained from other approximation methods regularly to simulation results obtained using Monte Carlo sampling.

#### Sigma point methods

The sigma point approach (Van Der Merwe, 2004) is one of the computationally most efficient methods to approximate a population of cells. Given ***n***_*b*_ random effects, we need to compute 2***n***_***b***_ + 1 (sigma) points and propagate them through the ODE. These points are usually located at 0 and ±***h*** along the parameter axes in each random effect direction, where ***h*** is a chosen step size, which is set to 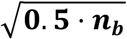. If ***∑*** is the unit matrix, this will yield ***b***^***i,SP***^ = ***he***_***i***_ with ***e***_***i***_ being the ***i***-th unit vector. In the general case, the sigma points get transformed with the lower Cholesky decomposition of the covariance matrix:

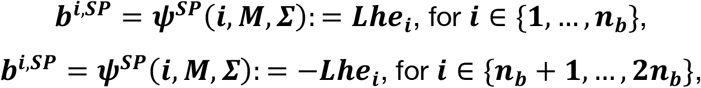

and

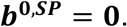

Additionally, the sigma points carry weights 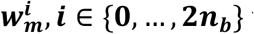 for the means and weights 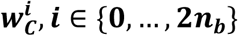 for the covariances. They are computed as:

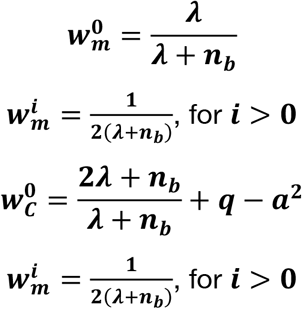

where we fixed ***a*** = **0.7** (it should hold: **0** < ***a*** ≤ **1**), ***q*** = **2** (it should hold ***q*** ≥ **1**) and **λ** = ***n***_***b***_(***a*^2^ − 1**). These weights are then used to compute means, variances and covariances of observables.

#### Dirac mixture models

The sigma point approximation may be computationally efficient, but it reflects the mean and variance only for linear systems exactly. Therefore, it can be inaccurate for highly nonlinear applications such as nonlinear ODE systems. Since in our case, massive Monte Carlo sampling was computationally infeasible and sigma points approximations provided poor results, we decided to use Dirac mixture distribution (DMD) approximations (Gilitschenski & Hanebeck, 2013). A DMD is a small set of points, which attempts to accurately reflect a multivariate normal distribution by fulfilling an optimality condition on the approximation quality. The number of these Dirac points can be freely chosen, in order to adapt the approximation quality. Thus, a DMD can either be interpreted as a small, but optimal sample, or as a sigma point method with adjustable accuracy. This allowed us to balance computational effort and numerical accuracy with high flexibility. However, this has the disadvantage that the DMD with the desired properties must be computed before it can be used for parameter estimation. To do so, a multivariate standard normal distribution with the desired dimension ***n***_***b***_ is approximated by a mixture of ***n***_***D***_ Dirac-delta distributions. We decided to use ***n***_***D***_ = 42, since this yields ***n***_***D***_/***n***_***b***_ = **6**, which are three times as many points as in the case of the sigma point approximation, which has ***n***_***D***_ = **2*n***_***b***_ + **1**. This guarantees a substantially higher approximation accuracy than the sigma point approximation.

The DMD for the smaller models (***n***_***b***_ = **6** and ***n***_***b***_ = **4**) were computed from the largest one by integrating out the corresponding columns. In this way, we could make sure that the calibrated models are really nested and can be analyzed with means such as the BIC. Otherwise, slightly different approximation accuracies due to different numbers of mixture points can dominate the parameter estimation results and may lead to incorrect conclusions when applying model criteria such as the BIC.

In order to compute the locations of the ***n***_***D***_ Dirac-points 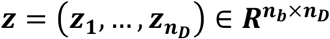, a modified Cramér-von Mises-distance between distributions is defined based on a measure called localized cumulative distribution (LCD) (Hanebeck & Klumpp, 2008). The LCD is a substitute for the cumulative distribution for multivariate probability density functions. For an ***n***_***b***_-dimensional probability density function 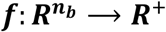, the corresponding LCD is defined as

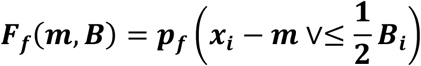

with 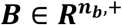, 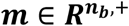 and ***p***_***f***_ being the probability given the density function ***f***. This is used to define a (so-called modified Cramér-von Mises) distance between two densities ***f*** and ***g*** in the following way:

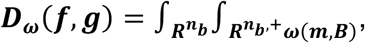

where an additional weighting function *ω* can be used.

We chose ***f*** to be a multivariate standard normal distribution ***N***(0, 1) with dimension ***n***_***b***_ = 7, and ***g*** to be a ***DMD***(***z***) with ***n***_***D***_ = 42 points at the locations ***z***^***i***^, ***i*** = **1**, …, ***n***_***D***_ and the weighting to

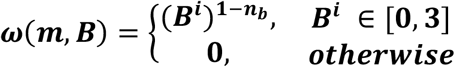

Then we optimized these locations in order to minimize the distance ***D*** and obtained the Dirac points for a standard normal distribution:

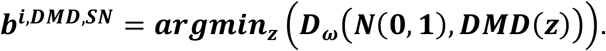

Once the DMD points ***b***^***i,DMD,SN***^ were computed, they can be transformed with the lower Cholesky decomposition ***L*** of the covariance matrix ***∑***

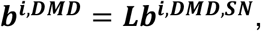

and all computations are carried out in the same way as for Monte Carlo sampling.

### Parametrizations of covariance matrices

To parametrize the covariance matrices *∑* of the random effects, we tested different approaches, as described in Pinheiro and Bates (1996):

- A log-matrix parametrization, in which a symmetric matrix *R* is parametrized, which fulfils *∑* = *exp*(*R*).
- The Givens parametrization, in which the eigenvalues are parametrized logarithmically as a diagonal matrix 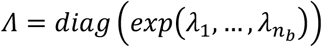 and the system of eigenvectors is parametrized as a product of *n*_*u*_ = *n*_*b*_ (*n*_*b*_ − 1)/2 rotations 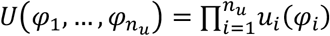, where the *φ*_*i*_ ∈ [0, *π*] are rotation angles. This yields *∑* = *U*^*T*^*∧U*.

Since none of these methods led to satisfactory results, we designed a novel approach based on Lie-theoretic considerations: We parametrized the eigenvalues logarithmically as diagonal matrix

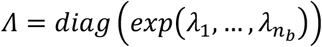

and defined

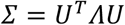

with *U* being a rotational matrix. However, we parametrized *U* not directly, as done in the Givens-formula, but indirectly, via the Lie algebra of the rotation group *SO*(*n*), i.e., we set

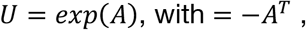

which we assume to have entries *A*_*i,j*_ ∈ [−*b*,*b*], with *b* ≫ 1, where *b* is an arbitrarily chosen bound, for which we used *b* = 20. Thus, we only need to parametrize the antisymmetric matrix *A*, which is a much simpler task than parametrizing the rotations.

This approach takes advantage of the fact that the exponential map is a local diffeomorphism around the unit elements of the Lie group and the Lie algebra. Furthermore, *A* = 0 being a natural initial guess for the optimization problem, lies at the center of the parameter interval. Compared to the Givens parametrization, this has the advantage of reducing the dependence of the *φ*_*i*_ on each other and hence improving the optimizer convergence.

### Parameter estimation for the single-cell models

The parameter estimation for the single-cell models was carried out using the MATLAB-based toolbox PESTO (Stapor et al, 2018). Model calibration was performed using the trust-region-reflective algorithm of the function fmincon (MATLAB Release 2017a), which is part of the MATLAB optimization toolbox. The optimization algorithm was provided with gradients and the Fisher information matrix from forward sensitivity analysis. The mixed-effect model, the different methods for parametrizing the covariance matrix of random effects, and the corresponding sensitivities were constructed using the MATLAB-based toolbox MEMOIR (Fröhlich et al, 2018). MEMOIR also assembled the objective function, which was passed to PESTO, as the log-posterior function of observing a parameter vector given the experimental data, assuming a Gaussian noise model for the population average data, and the mean values, variances and covariances of the single-cell snapshot data. The numerical integration of the ODE systems and their forward sensitivities was carried out using the C++ and MATLAB-based toolbox AMICI (Fröhlich et al, 2017), which interfaces the CVODES solver from the SUNDIALS suite (Hindmarsh et al, 2005).

## QUANTIFICATION AND STATISTICAL ANALYSIS

### Model selection criteria

Model selection for the single-cell models was based on the Bayesian information criterion (BIC) to have a asymptotically consistent model selection criterion. The BIC accounts for the maximum log-likelihood of each model (as a measure for fit quality) and model complexity. The BIC is computed as follows:

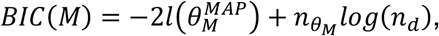

where *M* is the model, *l* log-posterior (or the log-likelihood) with the maximum a posteriori (or maximum likelihood) estimate *θ*^*MAP*^, 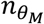 is the number of parameters of *M* and *n*_*d*_ is the number of data points. Especially for models with many data points (like in our case), the BIC penalizes model complexity stronger than the Akaike information criterion (AIC) and may hence give stronger indications for model selection.

### Translocation rates and compartmental volumes

Our single-cell model selection reveals the import and export rates to be variable, and it predicts the heterogeneity of the import rates to be higher than the heterogeneity of the export rates (Figure 6A): In order to quantify this difference more precisely, we used the estimated decadic translocation rate constants *Import*_*STATS*_ and *Export*_*STATS*_ and the estimated covariance matrix of random effects *∑* for the simulation of 1000 cell population with 100 000 cells each. In this way, we can compute the ratios of the coefficients of variation *CV* of the rate constants in linear scale for each cell population. This yields for the respective mean values

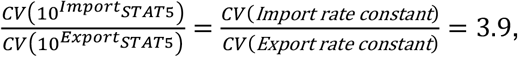

and a value of 1.6 for the standard deviation of this ratio.

The inverses of the import and export rates on the other hand are proportional to the expected time for a molecule of STAT5 to be imported or exported, respectively (having the unit s/mol or only s, if we consider molecules instead of abundances):

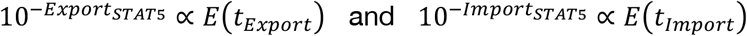

If we assume that molecular transport is driven by diffusion, this expected time for a molecule to be exported should depend on the radius of the cell’s nucleus *r*_*nuc*_, which is related to the nucleic volume by 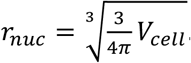. On the other hand, we assume the expected time for a molecule to be imported to depend on the thickness of the cytoplasmic shell *t*_*cyt*_ around the nucleus, which is related to the compartmental volume via 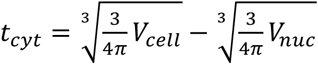 (see Figure 6B).

Results from the theory of cellular diffusion processes (Bressloff, 2014) state that relations

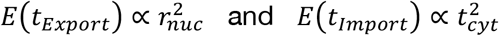

should hold. We confirmed the first relation by simulating 10 000 random walks in three dimensions with for different nucleic radii (Figure 6C). In such a random walk, which we always started in the center of the nucleus, a molecule could move along each coordinate axis either forward or backward with a fixed step size until it reached the boundary of the nucleic sphere. The mean of the computation times of these random walks was computed and used as a proxy for the expected export time of a STAT5 molecule. For the nuclear import process, we assumed that that *r*_*cyt*_ ≪ *r*_*nuc*_, which allowed us to approximate the import process by a random walk between two planes with distance *t*_*cyt*_ in three dimensions. We simulated again 10 000 random walks, which were constraint to stay between the planes, starting at the first one (the cell membrane) and stopped them when the second one (the nucleic membrane) was reached. By carrying out the corresponding computations for different distances between the planes, we also confirmed the second relation (Figure 6C).

Based on these considerations, we conclude that the relation

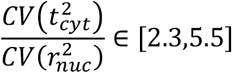

should hold. Evaluating the results from confocal fluorescence microscopy z-stack images, the corresponding coefficients of variation indeed yielded the ratio

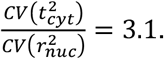

### Inference of the survival criterion

Similar to the work by Bachmann et al. (2011), we compared different criteria for survival against each other. Considered criteria, whether a single-cell would survive, were

- the total amount of pSTAT5 in the nucleus, which the respective cell is exposed to averaged over the time interval [0,***t***_***end***_].
- the maximal ratio of pSTAT5 in the nucleus over the total amount of STAT5, which the respective cell is exposed to averaged over the time interval [0,***t***_***end***_].

We checked these two proposed criteria for ***t***_***end***_ from 5 to 180 min, in steps of 5 min. The necessary thresholds for the respective survival signals were fitted to the survival data based on the selected single-cell model at the optimal parameter value. Since we observe survival also in the absence of Epo, we additionally fitted an offset for this basal survival rate.

The BIC values for these criteria and thresholds yielded that the criteria based on the ratio of pSTAT5 with a threshold ***t***_***end***_ = 180 min was the preferred model for cell survival. The BIC values for the two criteria are shown in Supplemental Figure S7A.

