## Supplementary file for "Cell-to-cell variability in JAK2/STAT5 pathway components and cytoplasmic volumes define survival threshold in erythroid progenitor cells"

**Authors/Affiliations**

Lorenz Adlung[1,8], Paul Stapor[2,3,8], Christian Tönsing[4,5,8], Leonard Schmiester[2,3], Luisa E. Schwarzmüller[1], Dantong Wang[2,3], Jens Timmer[4,5,*] , Ursula Klingmüller[1,6,*], Jan Hasenauer[2,3,7,*], Marcel Schilling[1,9,*]

[1] Division Systems Biology of Signal Transduction, German Cancer Research Center (DKFZ), 69120 Heidelberg, Germany

[2] Helmholtz Zentrum München - German Research Center for Environmental Health, Institute of Computational Biology, 85764 Neuherberg, Germany

[3] Technische Universität München, Center for Mathematics, Chair of Mathematical Modeling of Biological Systems, 85748 Garching, Germany

[4] Institute of Physics, University of Freiburg, 79104 Freiburg, Germany

[5] CIBSS – Centre for Integrative Biological Signalling Studies, University of Freiburg, 79104 Freiburg, Germany

[6] Translational Lung Research Center (TLRC), Member of the German Center for Lung Research (DZL), 69120 Heidelberg, Germany

[7] Faculty of Mathematics and Natural Sciences, University of Bonn, 53113 Bonn, Germany

[8] These authors contributed equally

[9] Lead Contact

### 1 Supplemental Figures

#### 1.1 Figure S1

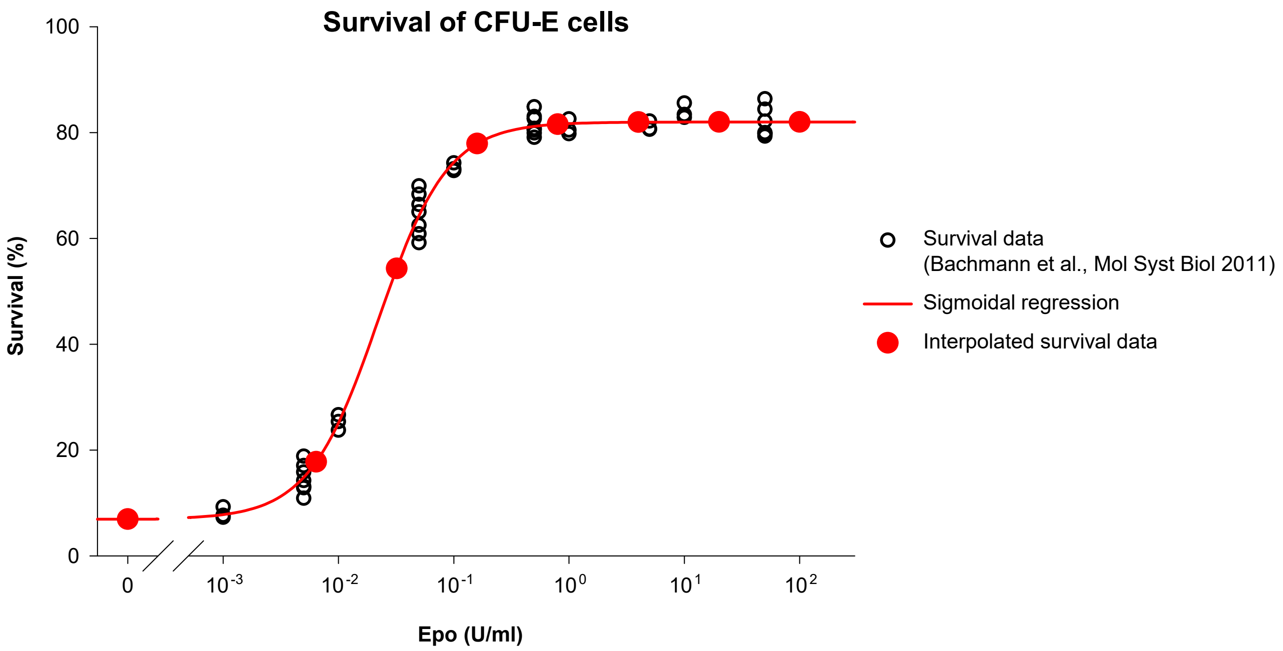

**Figure S1 (related to Figure 1):** Survival data for CFU-E cells. The percentage of surviving CFU-E cells cultured 24 h in various Epo concentration measured by TUNEL assay as reported in Bachmann et al., Mol Syst Biol 2011 are shown as open black circles. A sigmoidal regression based on a four parameter Hill function was used (solid red line) to interpolate survival data for the Epo concentrations used in Figure 1 (solid red circles).

#### 1.2 Figure S2

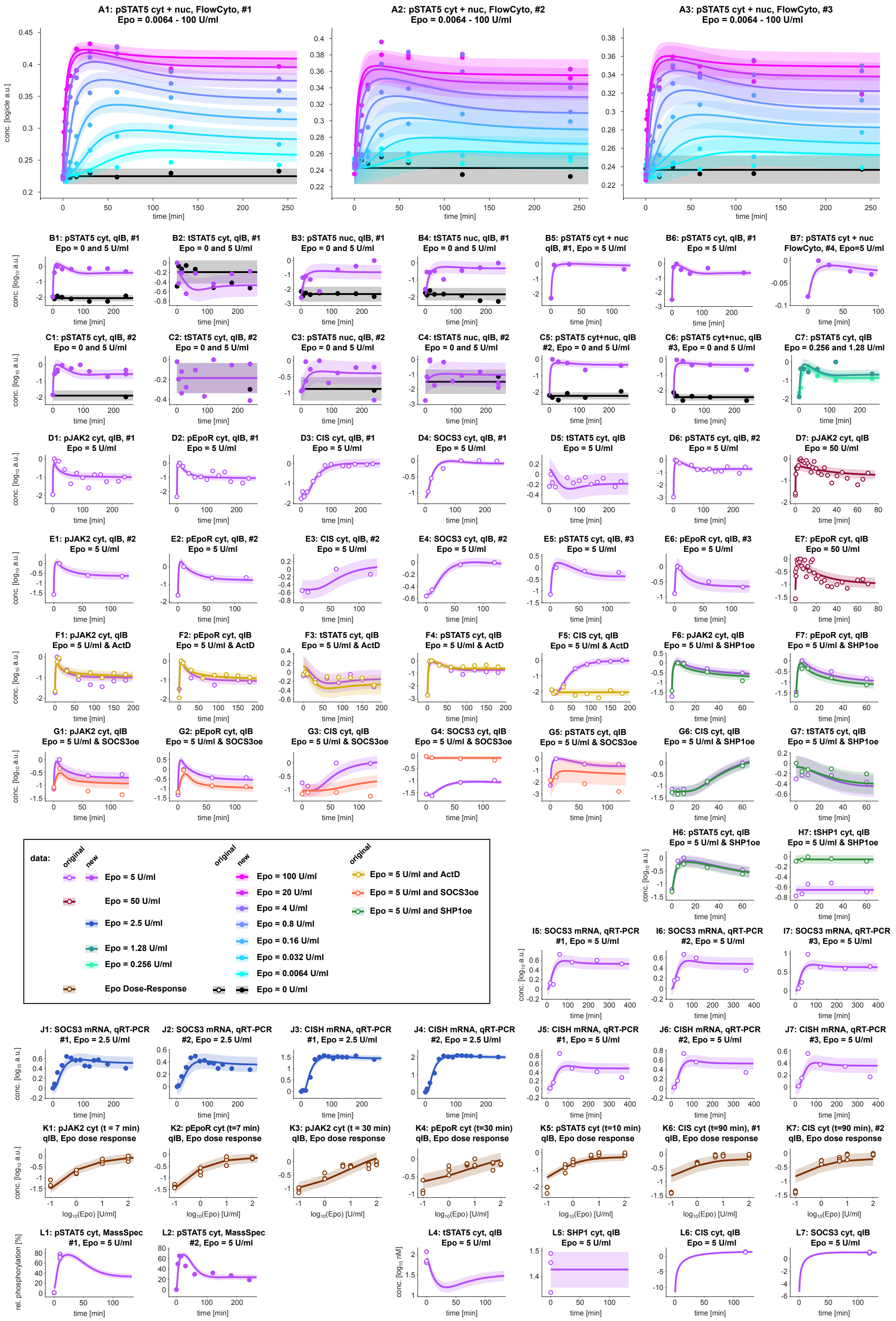
A1: pSTAT5 cyt + nuc, FlowCyto, #1
Epo = 0.0064 - 100 U/ml
A2: pSTAT5 cyt + nuc, FlowCyto, #2
Epo = 0.0064 - 100 U/ml
A3: pSTAT5 cyt + nuc, FlowCyto, #3
Epo = 0.0064 - 100 U/ml
conc. [logicle a.u.]
time [min]
B1: pSTAT5 cyt, qIB, #1
Epo = 0 and 5 U/ml
B2: tSTAT5 cyt, qIB, #1
Epo = 0 and 5 U/ml
B3: pSTAT5 nuc, qIB, #1
Epo = 0 and 5 U/ml
B4: tSTAT5 nuc, qIB, #1
Epo = 0 and 5 U/ml
B5: pSTAT5 cyt + nuc
qIB, #1, Epo = 5 U/ml
B6: pSTAT5 cyt, qIB, #1
Epo = 5 U/ml
B7: pSTAT5 cyt + nuc
FlowCyto, #4, Epo=5 U/ml
C1: pSTAT5 cyt, qIB, #2
Epo = 0 and 5 U/ml
C2: tSTAT5 cyt, qIB, #2
Epo = 0 and 5 U/ml
C3: pSTAT5 nuc, qIB, #2
Epo = 0 and 5 U/ml
C4: tSTAT5 nuc, qIB, #2
Epo = 0 and 5 U/ml
C5: pSTAT5 cyt+nuc, qIB
#2, Epo = 0 and 5 U/ml
C6: pSTAT5 cyt+nuc, qIB
#3, Epo = 0 and 5 U/ml
C7: pSTAT5 cyt, qIB
Epo = 0.256 and 1.28 U/ml
D1: pJAK2 cyt, qIB, #1
Epo = 5 U/ml
D2: pEpoR cyt, qIB, #1
Epo = 5 U/ml
D3: CIS cyt, qIB, #1
Epo = 5 U/ml
D4: SOCS3 cyt, qIB, #1
Epo = 5 U/ml
D5: tSTAT5 cyt, qIB
Epo = 5 U/ml
D6: pSTAT5 cyt, qIB, #2
Epo = 5 U/ml
D7: pJAK2 cyt, qIB
Epo = 50 U/ml
E1: pJAK2 cyt, qIB, #2
Epo = 5 U/ml
E2: pEpoR cyt, qIB, #2
Epo = 5 U/ml
E3: CIS cyt, qIB, #2
Epo = 5 U/ml
E4: SOCS3 cyt, qIB, #2
Epo = 5 U/ml
E5: pSTAT5 cyt, qIB, #3
Epo = 5 U/ml
E6: pEpoR cyt, qIB, #3
Epo = 5 U/ml
E7: pEpoR cyt, qIB
Epo = 50 U/ml
F1: pJAK2 cyt, qIB
Epo = 5 U/ml & ActD
F2: pEpoR cyt, qIB
Epo = 5 U/ml & ActD
F3: tSTAT5 cyt, qIB
Epo = 5 U/ml & ActD
F4: pSTAT5 cyt, qIB
Epo = 5 U/ml & ActD
F5: CIS cyt, qIB
Epo = 5 U/ml & ActD
F6: pJAK2 cyt, qIB
Epo = 5 U/ml & SHP1oe
F7: pEpoR cyt, qIB
Epo = 5 U/ml & SHP1oe
G1: pJAK2 cyt, qIB
Epo = 5 U/ml & SOCS3oe
G2: pEpoR cyt, qIB
Epo = 5 U/ml & SOCS3oe
G3: CIS cyt, qIB
Epo = 5 U/ml & SOCS3oe
G4: SOCS3 cyt, qIB
Epo = 5 U/ml & SOCS3oe
G5: pSTAT5 cyt, qIB
Epo = 5 U/ml & SOCS3oe
G6: CIS cyt, qIB
Epo = 5 U/ml & SHP1oe
G7: tSTAT5 cyt, qIB
Epo = 5 U/ml & SHP1oe
H6: pSTAT5 cyt, qIB
Epo = 5 U/ml & SHP1oe
H7: tSHP1 cyt, qIB
Epo = 5 U/ml & SHP1oe
data:
original
new
Epo = 5 U/ml
Epo = 50 U/ml
Epo = 2.5 U/ml
Epo = 1.28 U/ml
Epo = 0.256 U/ml
Epo Dose-Response
Epo = 100 U/ml
Epo = 20 U/ml
Epo = 4 U/ml
Epo = 0.8 U/ml
Epo = 0.16 U/ml
Epo = 0.032 U/ml
Epo = 0.0064 U/ml
Epo = 0 U/ml
Epo = 5 U/ml and ActD
Epo = 5 U/ml and SOCS3oe
Epo = 5 U/ml and SHP1oe
I5: SOCS3 mRNA, qRT-PCR
#1, Epo = 5 U/ml
I6: SOCS3 mRNA, qRT-PCR
#2, Epo = 5 U/ml
I7: SOCS3 mRNA, qRT-PCR
#3, Epo = 5 U/ml
J1: SOCS3 mRNA, qRT-PCR
#1, Epo = 2.5 U/ml
J2: SOCS3 mRNA, qRT-PCR
#2, Epo = 2.5 U/ml
J3: CISH mRNA, qRT-PCR
#1, Epo = 2.5 U/ml
J4: CISH mRNA, qRT-PCR
#2, Epo = 2.5 U/ml
J5: CISH mRNA, qRT-PCR
#1, Epo = 5 U/ml
J6: CISH mRNA, qRT-PCR
#2, Epo = 5 U/ml
J7: CISH mRNA, qRT-PCR
#3, Epo = 5 U/ml
K1: pJAK2 cyt (t = 7 min)
qIB, Epo dose response
K2: pEpoR cyt (t=7 min)
qIB, Epo dose response
K3: pJAK2 cyt (t = 30 min)
qIB, Epo dose response
K4: pEpoR cyt (t=30 min)
qIB, Epo dose response
K5: pSTAT5 cyt (t=10 min)
qIB, Epo dose response
K6: CIS cyt (t=90 min), #1
qIB, Epo dose response
K7: CIS cyt (t=90 min), #2
qIB, Epo dose response
L1: pSTAT5 cyt, MassSpec
#1, Epo = 5 U/ml
L2: pSTAT5 cyt, MassSpec
#2, Epo = 5 U/ml
L4: tSTAT5 cyt, qIB
Epo = 5 U/ml
L5: SHP1 cyt, qIB
Epo = 5 U/ml
L6: CIS cyt, qIB
Epo = 5 U/ml
L7: SOCS3 cyt, qIB
Epo = 5 U/ml
rel. phosphorylation [%]

**Figure S2 (related to Figure 2):** Complete data set and model output of population-average model dynamics. Empty circles indicate original data sets from Bachmann et al., Mol Syst Biol 2011, filled circles is new experimental data. All data shown enter the analyses. Model simulations are shown as solid lines in the color of the corresponding experimental conditions. Shading depicts the error model. Measured entities, experimentation techniques, replicate number (rep #) and experimental conditions as indicated in the subplot titles. cyt: only cytopasmatic fraction, nuc: only nuclear fraction, cyt + nuc: whole cell lysates. FlowCyto: Flow cytometry (A1-3, B7), qIB: quantitative immunoblotting (B1-H7, L4-L7), qRT-PCR: quantitative Real-Time Transcription PCR (I5-7, J1-7), MassSpec: Mass spectrometry (L1-L2)

#### 1.3 Figure S3

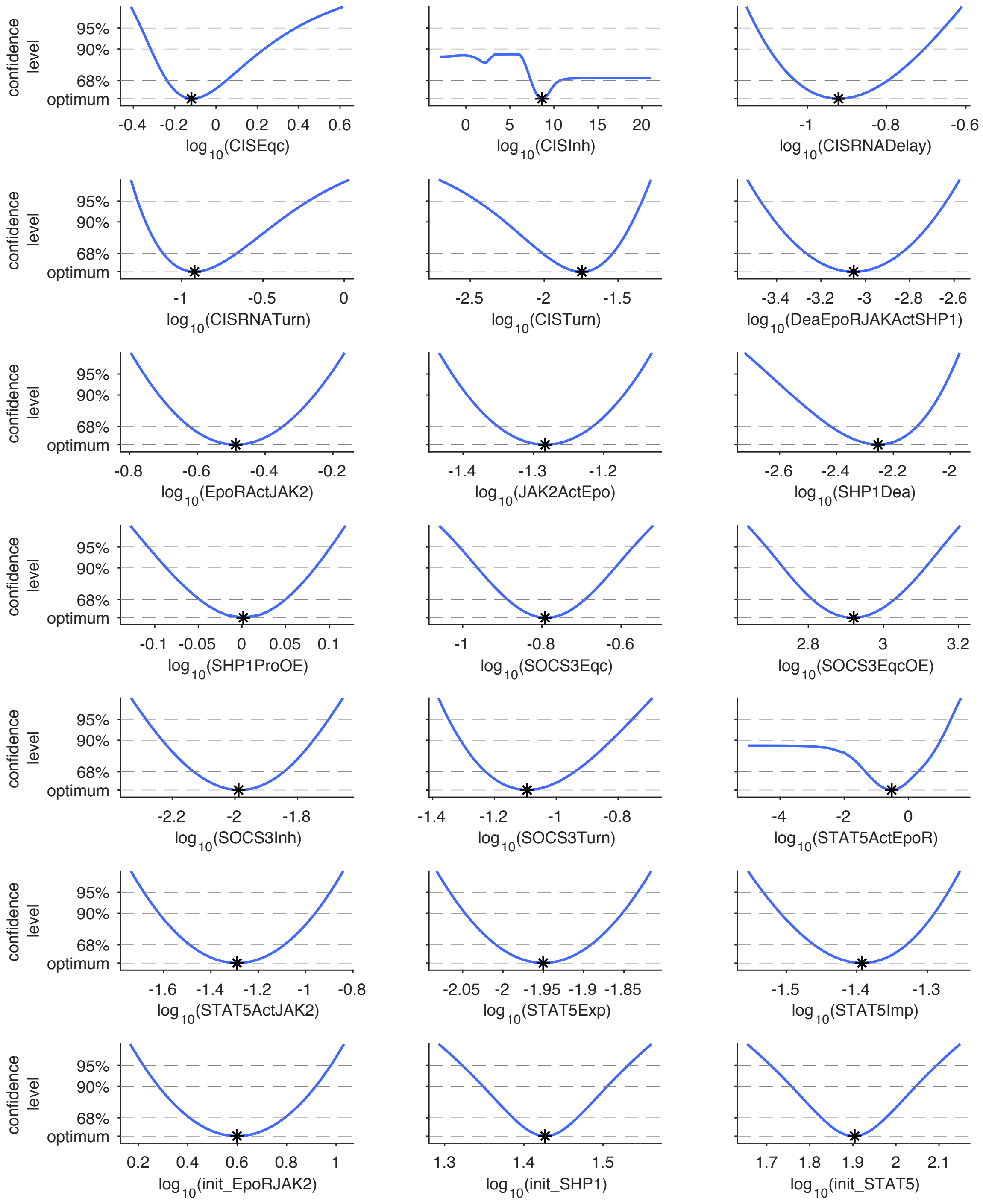
confidence level
95%
90%
68%
optimum
-0.4 -0.2 0 0.2 0.4 0.6
log 10(CISEqc)
0 5 10 15 20
log 10(CISInh)
-1 -0.8 -0.6
log 10(CISRNADelay)
-1 -0.5 0
log 10(CISRNATurn)
-2.5 -2 -1.5
log 10(CISTurn)
-3.4 -3.2 -3 -2.8 -2.6
log 10(DeaEpoRJAKActSHP1)
-0.8 -0.6 -0.4 -0.2
log 10(EpoRActJAK2)
-1.4 -1.3 -1.2
log 10(JAK2ActEpo)
-2.6 -2.4 -2.2 -2
log 10(SHP1Dea)
-0.1 -0.05 0 0.05 0.1
log 10(SHP1ProOE)
-1 -0.8 -0.6
log 10(SOCS3Eqc)
2.8 3 3.2
log 10(SOCS3EqcOE)
-2.2 -2 -1.8
log 10(SOCS3Inh)
-1.4 -1.2 -1 -0.8
log 10(SOCS3Turn)
-4 -2 0
log 10(STAT5ActEpoR)
-1.6 -1.4 -1.2 -1 -0.8
log 10(STAT5ActJAK2)
-2.05 -2 -1.95 -1.9 -1.85
log 10(STAT5Exp)
-1.5 -1.4 -1.3
log 10(STAT5Imp)
0.2 0.4 0.6 0.8 1
log 10(init_EpoRJAK2)
1.3 1.4 1.5
log 10(init_SHP1)
1.7 1.8 1.9 2 2.1
log 10(init_STAT5)

**Figure S3 (related to Figure 2):** Likelihood profiles of the pathway model with population-level data for all dynamic model parameters (blue lines). Black asterisks indicate the best fit and global optimum. Intersections of the likelihood profiles (blue solid lines) with the statistical thresholds for different confidence levels (gray dashed horizontal lines) indicate the profile likelihood-based confidence intervals. All parameters are identifiably with confidence levels of 68%. Only parameters CISInh and STAT5ActEpoR reveal practical non-identifiabilities for higher confidence levels. These non-identifiabilities could be resolved only by further measurements but they do not weaken the predictive power of the model as a unique optimum of the model parameters is found.

#### 1.4 Figure S4

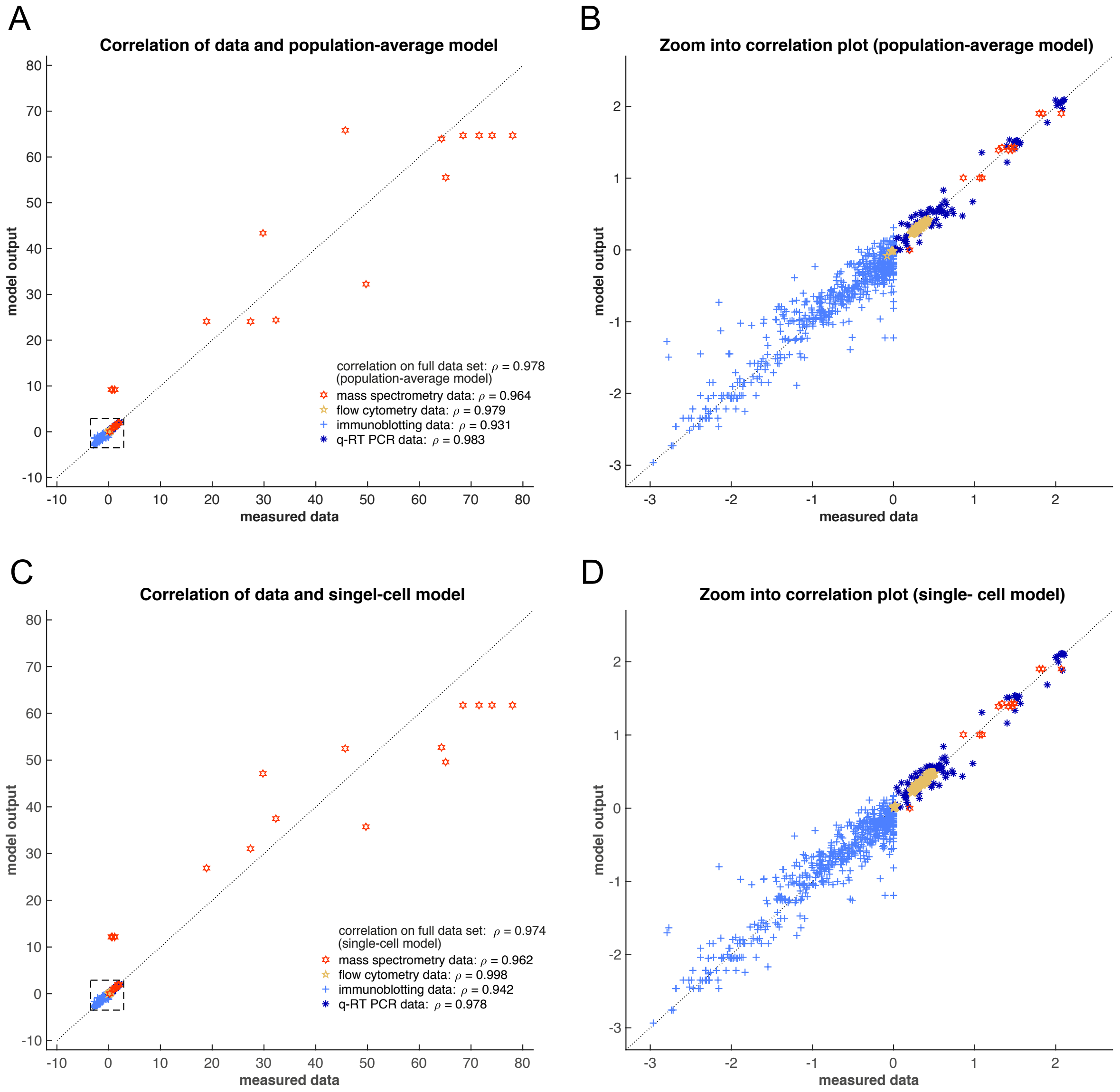

**Figure S4 (related to Figure 4):** Correlation of experimental data and model output. **(A)** Full data set for population-average model. Red hexagrams: mass spectrometry data (non-normalized, linear scale), yellow pentagrams: flow cytometry data (non-normalized, logicle scale), light blue plus signs: immunoblotting data (normalized, log-scale), deep blue asterisks: q-RT PCR data (non-normalized, log-scale). **(B)** Zoom into correlation plot for population-average model. **(C)** Full data set for single-cell model. **(D)** Zoom into correlation plot for single-cell model.

#### 1.5 Figure S5

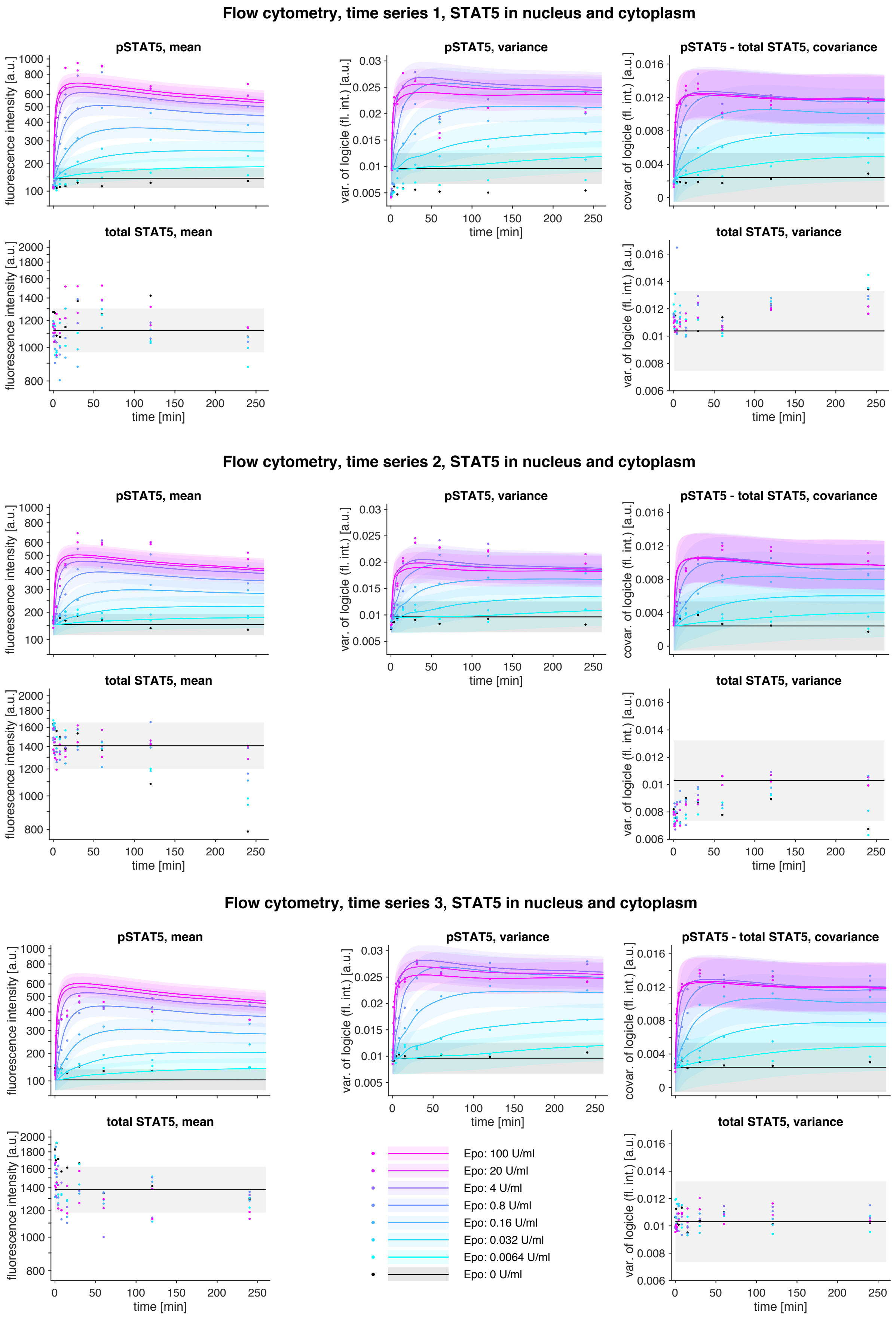

Flow cytometry, time series 1, STAT5 in nucleus and cytoplasm
pSTAT5, mean
fluorescence intensity [a.u.]
pSTAT5, variance
var. of logicle (fl. int.) [a.u.]
pSTAT5 - total STAT5, covariance
covar. of logicle (fl. int.) [a.u.]
total STAT5, mean
fluorescence intensity [a.u.]
total STAT5, variance
var. of logicle (fl. int.) [a.u.]
time [min]
Flow cytometry, time series 2, STAT5 in nucleus and cytoplasm
pSTAT5, mean
pSTAT5, variance
pSTAT5 - total STAT5, covariance
total STAT5, mean
total STAT5, variance
time [min]
Flow cytometry, time series 3, STAT5 in nucleus and cytoplasm
pSTAT5, mean
pSTAT5, variance
pSTAT5 - total STAT5, covariance
total STAT5, mean
total STAT5, variance
time [min]
Epo: 100 U/ml
Epo: 20 U/ml
Epo: 4 U/ml
Epo: 0.8 U/ml
Epo: 0.16 U/ml
Epo: 0.032 U/ml
Epo: 0.0064 U/ml
Epo: 0 U/ml

**Figure S5 (related to Figure 5):** Flow-cytometry data and model fit for the selected single-cell model. Three data sets were measured for eight Epo doses (colors), using flow cytometry and time-resolved experiments. Each experiment shows time-courses of population means (left) of pSTAT5 (upper panel) and total STAT5 (lower panel), and time-course variances and covariance of both species plotted as correlation matrix trajectories (right).

#### 1.6 Figure S6

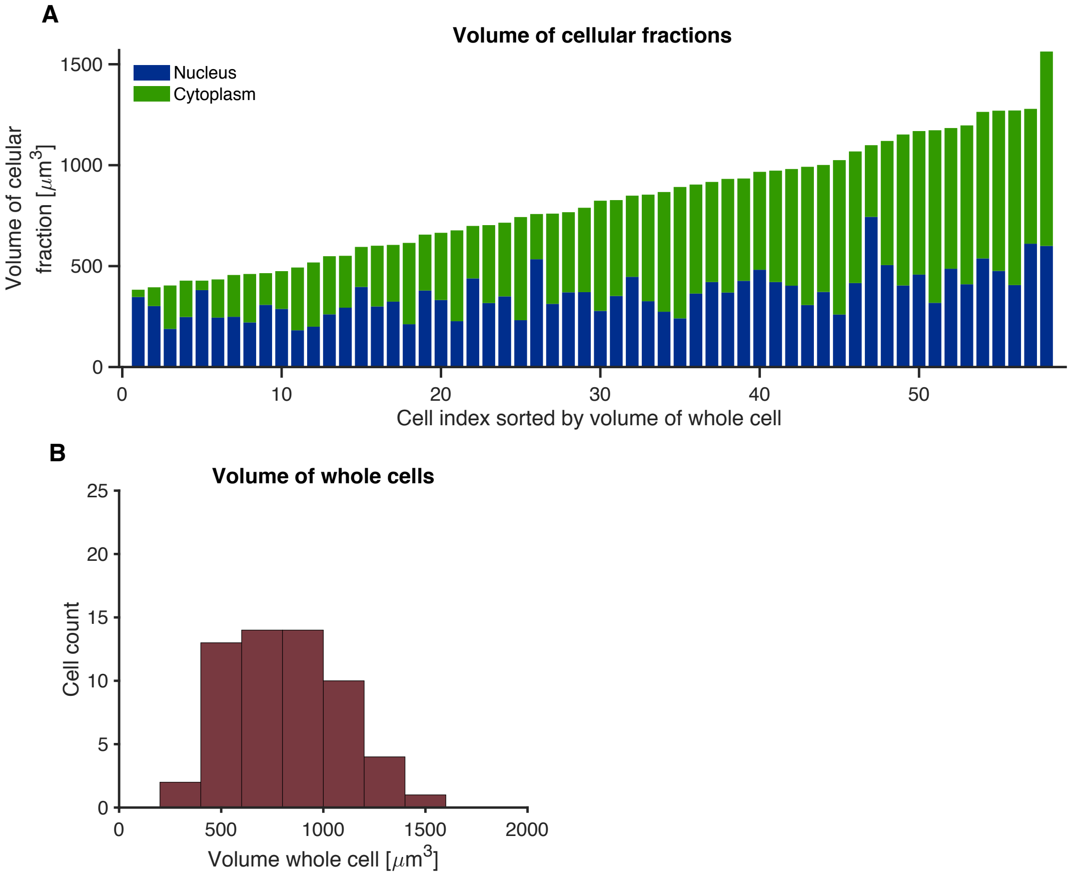

**Figure S6 (related to Figure 6):** Analysis of cytoplasmic and nuclear volumes from unstimulated growth factor-depleted CFU-E cells from by confocal microscopy and 3D reconstruction. **(A)** Measured nucleus and cytoplasm volumes from n = 58 cells. **(B)** Distribution of whole cell volumes.

#### 1.7 Figure S7

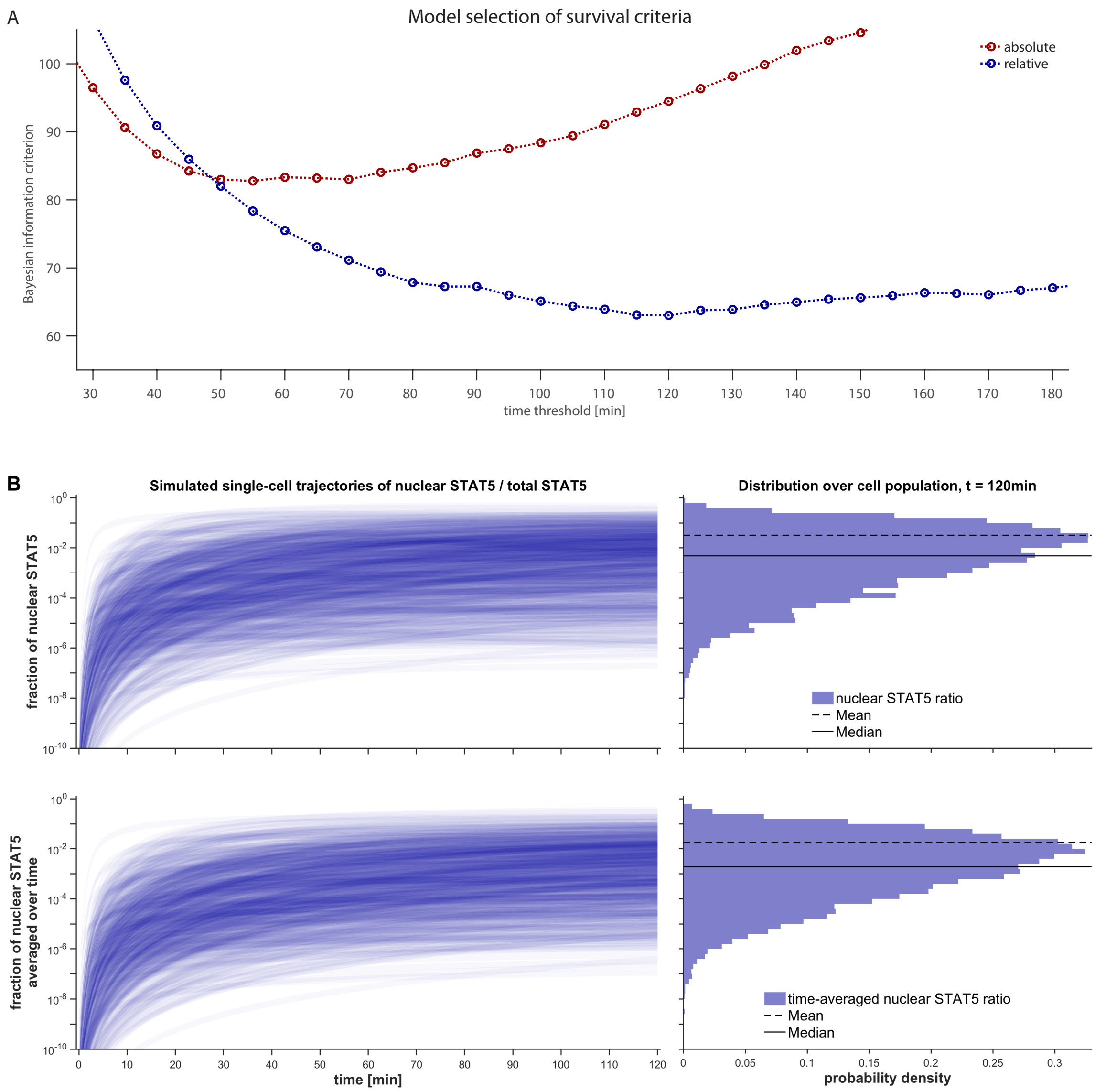

**Figure S7 (related to Figure 7): (A)** BIC results for model selection for different possible models of cell survival. pSTAT5 must be in the nucleus to ensure cell survival. An absolute criterion, i.e., the abundance of STAT5 in the nucleus (red) and a relative criterion, i.e., the percentage of a cell's STAT5 in the nucleus (blue), were compared against each other for different time intervals. Survival thresholds and offset parameters were fitted and the fit quality was compared. The relative criterion for a time interval of 120 minutes provided the best results. **(B)** Trajectories (left panel) for an in-silico population of 10 000 cells at $EC_{50}$-dose of Epo, which show the fraction of pSTAT5 in the cell's nucleus (upper panel) and the time-average of this fraction from $t = 0$ to the current time (lower panel). The right panel shows the histograms over the population for the two quantities at $t = 120\,\text{min}$, which demonstrate the log-normal distribution of the survival signal. The median in the lower right panel represents the survival threshold.
